# Graph-based pangenome reveals structural variation dynamics during cucumber breeding

**DOI:** 10.1101/2025.07.17.665377

**Authors:** Xuebo Zhao, Jingyin Yu, Jie Zhang, Honghe Sun, Shan Wu, Jiantao Zhao, Yao Zhou, Sue A. Hammar, Ying-Chen Lin, Zhonghua Zhang, Sanwen Huang, Ronald T. Dymerski, Feifan Chen, Yiqun Weng, Rebecca Grumet, Yong Xu, Zhangjun Fei

**Affiliations:** Boyce Thompson Institute, Ithaca, NY 14853, USA; State Key Laboratory of Vegetable Biobreeding, National Engineering Research Center for Vegetables, Beijing Key Laboratory of Vegetable Germplasms Improvement, Beijing Vegetable Research Center, Beijing Academy of Agriculture and Forestry Science, Beijing, 100097, China; Institute of Botany, Chinese Academy of Sciences, Beijing 100093, China; College of Life Sciences, University of Chinese Academy of Sciences, Beijing 100094, China; Department of Horticulture, Graduate Program in Plant Breeding, Genetics and Biotechnology, Michigan State University, East Lansing, MI 48824, USA; Engineering Laboratory of Genetic Improvement of Horticultural Crops of Shandong Province, College of Horticulture, Qingdao Agricultural University, Qingdao 266109, China; National Key Laboratory of Tropical Crop Breeding, Shenzhen Branch, Guangdong Laboratory of Lingnan Modern Agriculture, Genome Analysis Laboratory of the Ministry of Agriculture and Rural Affairs, Agricultural Genomics Institute at Shenzhen, Chinese Academy of Agricultural Sciences, Shenzhen, China; National Key Laboratory of Tropical Crop Breeding, Chinese Academy of Tropical Agricultural Sciences, Haikou, China; Department of Plant and Agroecosystem Sciences, University of Wisconsin, Madison, WI 53706, USA; USDA-ARS Vegetable Crops Research Unit, Madison, WI 53706, USA; Xianghu Laboratory, Hangzhou 311231, China; Syngenta Group China, Beijing, 100012, China

## Abstract

Structural variants (SVs) represent a significant yet underexplored component of plant genome diversity. Analyzing SV dynamics is important for understanding their contributions to phenotypic variation and their influence on genome evolution. Here, we present a graph-based pangenome for cucumber, constructed from 39 reference-level genome assemblies. The pangenome captures 171,892 high-confidence SVs and enables genotyping of these SVs across 443 cucumber accessions, representing diverse geographic origins and spanning wild progenitors, landraces, and modern cultivars. Our analyses reveal that during cucumber domestication, a substantial portion of mildly deleterious SNPs are retained, whereas SVs are consistently purged, highlighting their potentially highly deleterious nature. During the geographic expansion of cucumber, a reduction in SV burden and a younger age of SVs than SNPs were observed, suggesting stronger purifying selection acting on SVs. Additionally, notable gene flow from wild population to African and European populations was detected, resulting in an increased SV burden, potentially due to hitchhiking effects. Importantly, incorporating SV burden into genomic prediction models improved the prediction accuracy for several agronomically important traits. This study illuminates the dynamics of SVs during cucumber domestication and improvement, highlights the complex interplay between SVs and selection pressures, and underscores the extensive implications of SVs for future cucumber breeding.

## Introduction

Structural variants (SVs), including insertions, deletions, duplications, inversions, and translocations, represent a significant component of genomic diversity^1^. SVs have substantial impacts on genome structure and function and play substantial roles in phenotypic diversity and adaptive evolution. However, detecting and characterizing SVs has been challenging due to their size and complexity^2^. As a result, SVs remain less explored than single nucleotide polymorphisms (SNPs), particularly in terms of their population-level dynamics^3^. Understanding the distribution, dynamics, and impacts of SVs is important for advancing plant biology and improving economically important crops^4–6^. Recent advances in sequencing technologies and computational tools have enabled large-scale exploration of SVs, providing deeper insights into their evolutionary dynamics and functional impacts^7^. Notably, graph-based pangenomes have emerged as a powerful approach for accurately capturing SVs within complex genomes. By integrating genetic diversity of multiple accessions into a unified graph structure, this method allows for comprehensive identification of SVs across entire populations, offering a more complete and nuanced representation of genomic variation compared to traditional reference-based approaches^8^.

Cucumber (*Cucumis sativus* L.) is an economically important crop and also serves as a model system for studies on sex determination and plant vascular biology^9^. Previous population genomic studies utilizing short-read sequencing have provided valuable insights into the domestication history and divergence of cucumber^10,11^. Recent chromosome-scale genome assemblies have revealed the type, size, and distribution of SVs in wild and cultivated cucumbers, offering a deeper understanding of cucumber karyotype evolution^12–14^. Despite these advancements, the population dynamics and biological effects of SVs in cucumber remain poorly understood. Investigating these aspects is important for elucidating the evolutionary forces shaping SVs and evaluating their potential in genome-wide association studies for trait discovery and variety improvement.

In this study, we present reference-quality genomes assembled from highly accurate PacBio HiFi reads for 27 wild and cultivated cucumber accessions. Combined with 12 previously reported chromosome-scale genome assemblies, we constructed a graph-based pangenome for cucumber, capturing a total of 171,892 high-confidence SVs. By leveraging the constructed graph-based pangenome and newly generated deep resequencing data, we genotyped these SVs in a diverse collection of 443 cucumber accessions, representing a broad range of geographic origins and encompassing wild progenitors, landraces, and modern cultivars. This comprehensive dataset enabled us to characterize the spatiotemporal spectrum of SV burden during cucumber domestication, geographic expansion, and introgression. Notably, incorporating SV burden improves genomic prediction accuracy for several agronomically important traits. Our findings provide valuable insights into the dynamics of SVs and highlight their extensive implications for cucumber breeding.

## Results

### Assembly and annotation of reference-quality genomes

To capture the genetic diversity of cucumber, we selected 27 accessions to generate reference-quality genome assemblies using PacBio HiFi long-read sequencing. This collection comprised five wild (*C. sativus* var. *hardwickii*), four Xishuangbanna (*C. sativus* var. *xishuangbannanesis*, cultivated in the tropical southwestern region of China and surrounding regions including Thailand, Laos, and Myanmar), and 18 cultivated (*C. sativus* var. *sativus*) accessions. Among the cultivated accessions, three were landraces primarily grown in India and 15 were cultivars from diverse geographic regions and market groups, including East Asia, Central/West Asia, Africa, Europe, and America (**Fig. 1a**). A total of 318.5 Gb of HiFi sequencing data were generated, with an average of 11.8 Gb (32.2×) per accession (**Supplementary Table 1**). Using these HiFi sequences, we successfully generated chromosome-scale assemblies for the 27 cucumber accessions. The final genome assembly sizes ranged from 259.1 Mb to 302.1 Mb, with an average of 286.8 Mb, and the contig N50 sizes ranged from 5.25 Mb to 29.81 Mb, with an average of 16.26 Mb (**Supplementary Table 1** and **Supplementary Fig. 1**). On average, 95.7% of the assembled contigs were anchored and ordered onto the seven cucumber chromosomes, with unanchored contigs primarily consisting of rDNA and other highly repetitive sequences. BUSCO^15^ assessment revealed an average completeness rate of 98.36% for these assemblies. Further evaluation using a k-mer-based approach implemented in Merqury^16^ indicated an average completeness rate of 97.9% and quality values (QV) all exceeding 60, equivalent to one base error per one million bases (**Supplementary Table 1**). Furthermore, the long terminal repeat (LTR) assembly index (LAI) scores ranged from 10.42 to 15.31, with an average of 13.38 (**Supplementary Table 1**), meeting the criteria for reference-level assemblies^17^. Collectively, these results confirmed the high quality of the 27 cucumber assemblies, which are superior or comparable to previously reported chromosome-level genome assemblies^12–14^. Additionally, a total of 21,347 to 22,361 protein-coding genes (with an average of 21,793) were predicted from these genome assemblies, and BUSCO completeness rates for the predicted genes ranged from 93.8% to 96.8% (with an average of 96.0%) (**Supplementary Table 1**). Repetitive sequences accounted for an average of 48.82% of the assemblies, ranging from 43.89% to 52.02% across the assembled genomes (**Supplementary Table 2**).

**Fig. 1.**
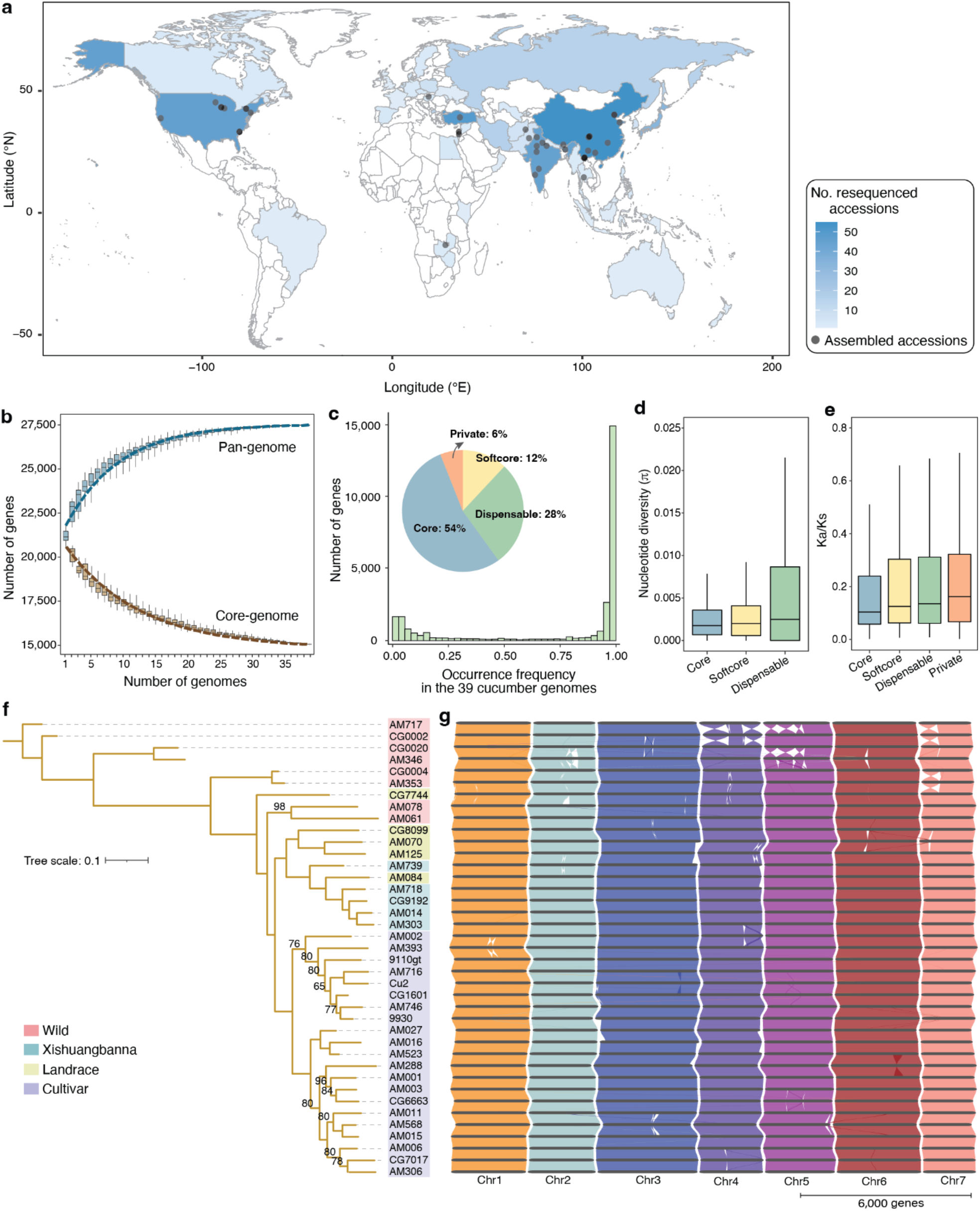
Pangenome of wild and cultivated cucumbers. **a**, Geographic distribution of assembled and resequenced cucumber accessions. The map was generated using the R package ggmap (https://cran.r-project.org/web/packages/ggmap/). **b**, Simulations of pangenome expansion and core-genome contraction. For each given number of accessions, 100 random combinations of accessions were sampled. **c**, Composition of the gene-based pangenome, categorized into core (present in all 39 accessions), softcore (present in more than 90% but not all accessions; 35 ≤ n < 39), dispensable (present in more than one but less than 90% of accessions; 2 ≤ n < 35), and private genes (present in only one accession). **d**, Nucleotide diversity (π) of core, softcore and dispensable genes. **e**, *Ka*/*Ks* values for core, softcore, dispensable and private genes. For each boxplot, the lower and upper bounds indicate the first and third quartiles, respectively, the center line indicates the median, and the whiskers extend to 1.5× the interquartile range. **f**, Phylogeny of the 39 cucumber accessions with reference-level genome assemblies. Bootstrap values below 100 are indicated on the branches. **g**, Synteny map across the 39 cucumber genomes, generated using GENESPACE (https://github.com/jtlovell/GENESPACE). The black segments represent chromosomes, while the colored blocks indicate regions of collinearity between genomes.

### Gene-based pangenome

To represent broader genetic diversity, we included the 27 genomes assembled in this study along with 12 previously reported chromosome-scale genome assemblies^13,14^ for gene-based pangenome construction (**Supplementary Fig. 2**). To mitigate potential issues caused by annotation artifacts, we reannotated the 12 previously reported genomes using the same pipeline applied to the 27 assembled genomes (**Supplementary Table 1**). Clustering the protein-coding genes from the 39 cucumber genome assemblies identified 27,779 distinct gene families (pan-gene clusters). The total number of pan-gene clusters plateaued when approximately 25 genomes were included, suggesting a near-saturation of pan-gene discovery within the scope of the analyzed accessions. (**Fig. 1b**). Across the 39 genomes, we identified 14,854 (53.47%) core, 3,324 (11.97%) softcore, 7,864 (28.31%) dispensable, and 1,737 (6.25%) private pan-gene clusters (**Fig. 1c**). Core genes exhibited lower nucleotide diversity (π) and lower nonsynonymous-to-synonymous substitution ratios (*K*a/*K*s) compared to dispensable and private genes (**Fig. 1d**,**e**). Moreover, core and softcore pan-gene clusters were enriched in functions fundamental to basic biological processes, such as transcription regulation, mRNA processing, and protein ubiquitination. In contrast, dispensable and private pan-gene clusters were enriched in functions related to enzyme activities and stress tolerance, such as enzyme metabolic processes, response to stimulus, and defense response (**Supplementary Fig. 3**).

**Fig. 2.**
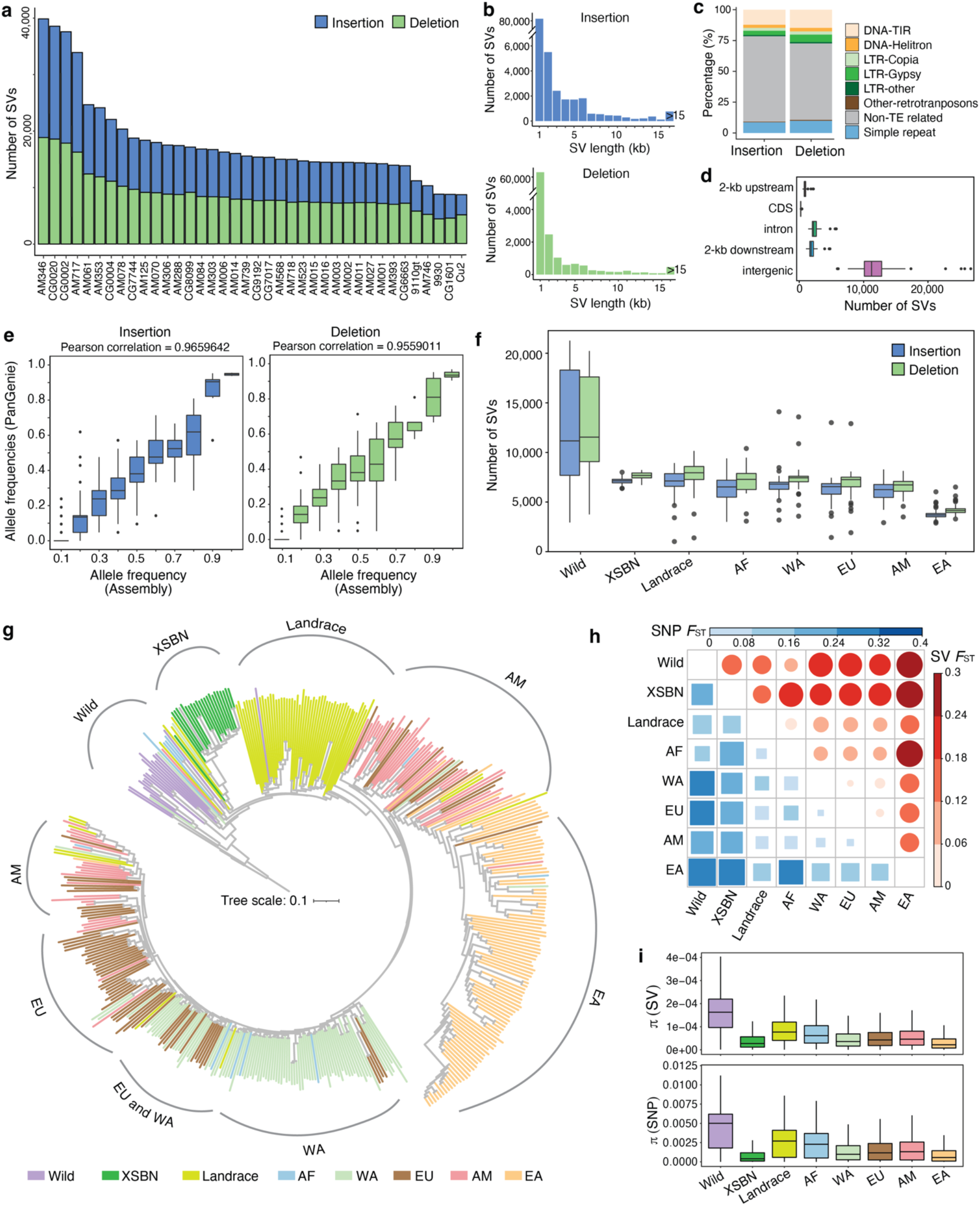
SV detection and genotyping in cucumber. **a**, Number of SVs in each accession. **b**, Length distribution of SVs. **c**, TE-and non-TE-derived SVs in the cucumber genome. **d**, Distribution of SVs in various genomic features. **e**, Comparison of allele frequencies derived from PanGenie genotyping of 10 accessions with those detected in the corresponding assemblies. **f**, Number of SVs identified in different cucumber populations. **g**, Phylogeny of the 443 cucumber accessions constructed using SVs. The curved arcs on the periphery indicate clade-level groupings, each labeled by the predominant population identity of its member accessions. **h**, Genome-wide fixation index (*F*_ST_) between cucumber populations based on SNPs and SVs. **i**, Nucleotide diversity of cucumber populations based on SNPs and SVs. For each boxplot, the lower and upper bounds indicate the first and third quartiles, respectively, the center line indicates the median, and the whiskers extend to 1.5× the interquartile range. XSBN, Xishuangbanna; AF, Africa; WA, Central/West Asia; EU, Europe; EA, East Asia; AM, America.

**Fig. 3.**
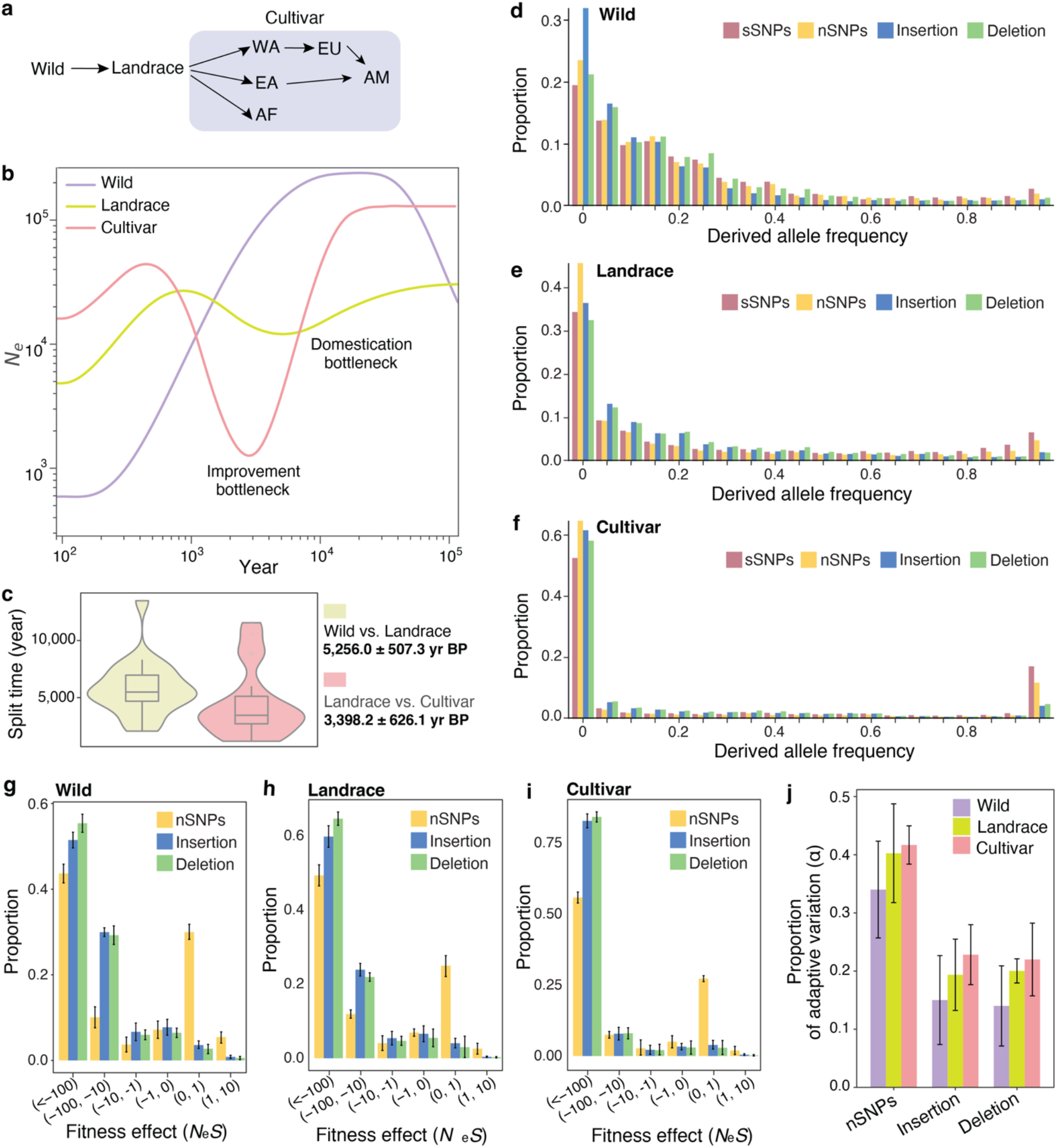
SV dynamics during cucumber domestication and improvement. **a**, Models of cucumber domestication and improvement processes. AF, Africa; WA, Central/West Asia; EU, Europe; EA, East Asia; AM, America. **b**, History of effective population sizes (*Ne*) in different cucumber populations. **c**, Distribution of estimated split times between wild cucumbers and landraces, and between landraces and cultivated cucumbers. Data are presented as violin plots with box plots. The bounds of the boxes represent the first and third quartiles, the center line indicates the median, and the whiskers extend to 1.5× the interquartile range. **d-f**, Site frequency spectrum (SFS) of synonymous SNPs (sSNPs), non-synonymous SNPs (nSNPs), insertions, and deletions in wild (**d**), landrace (**e**), and cultivated (**f**) cucumbers. **g-i**, Distribution of inferred fitness effects (*N*_*e*_*s*) for nSNPs, insertions, and deletions in wild (**g**), landrace (**h**), and cultivated (**i**) cucumbers. **j**, Proportion of adaptive variation (α) contributed by nSNPs and SVs in wild, landrace, and cultivated cucumbers. Values are derived from 20 bootstrap replicates.

Phylogenetic tree constructed for cucumber using 12,655 single-copy orthologous gene families indicated that cucumbers were first domesticated in India and later spread to nearby regions, including southwest China, where they formed the landraces and subsequently the Xishuangbanna group. Cucumber then spread globally, giving rise to various cultivars (**Fig. 1f**), consistent with findings from a previous resequencing study^10^. Further analysis revealed conserved macrosynteny across these genomes, whereas notable large-scale chromosomal rearrangements were observed across the genome, especially on chromosomes 4, 5, and 7 (**Fig. 1g**). These high-quality genome assemblies provide a robust platform for exploring genetic diversity and genotype-phenotype relationships in cucumber.

### Graph-based pangenome and SV genotyping

To comprehensively understand the evolutionary dynamics of SVs throughout cucumber breeding history, we constructed a graph-based pangenome using the 39 high-quality chromosome-scale assemblies, and identified 171,892 nonredundant SVs (≥20 bp), comprising 99,962 insertions and 71,930 deletions, spanning a total of 159,141,251 bp, with SVs in each genome contributing incrementally to the pangenome (**Fig. 2a, Supplementary Tables 3, 4** and **Supplementary Fig. 4**). Approximately 2.13% of the insertions and 1.79% of the deletions were greater than 10 kb, while 82.26% and 88.54% were shorter than 1 kb (**Fig. 2b** and **Supplementary Table 3**). A substantial portion (32.49%) of SVs were derived from transposable elements (TEs), with DNA transposons featuring terminal inverted repeats (DNA-TIR) being the most abundant type (**Fig. 2c** and **Supplementary Fig. 5**). About 5.09% overlapped with genic regions, affecting 19.5% of all protein-coding genes (**Fig. 2d** and **Supplementary Table 5**).

**Fig. 4.**
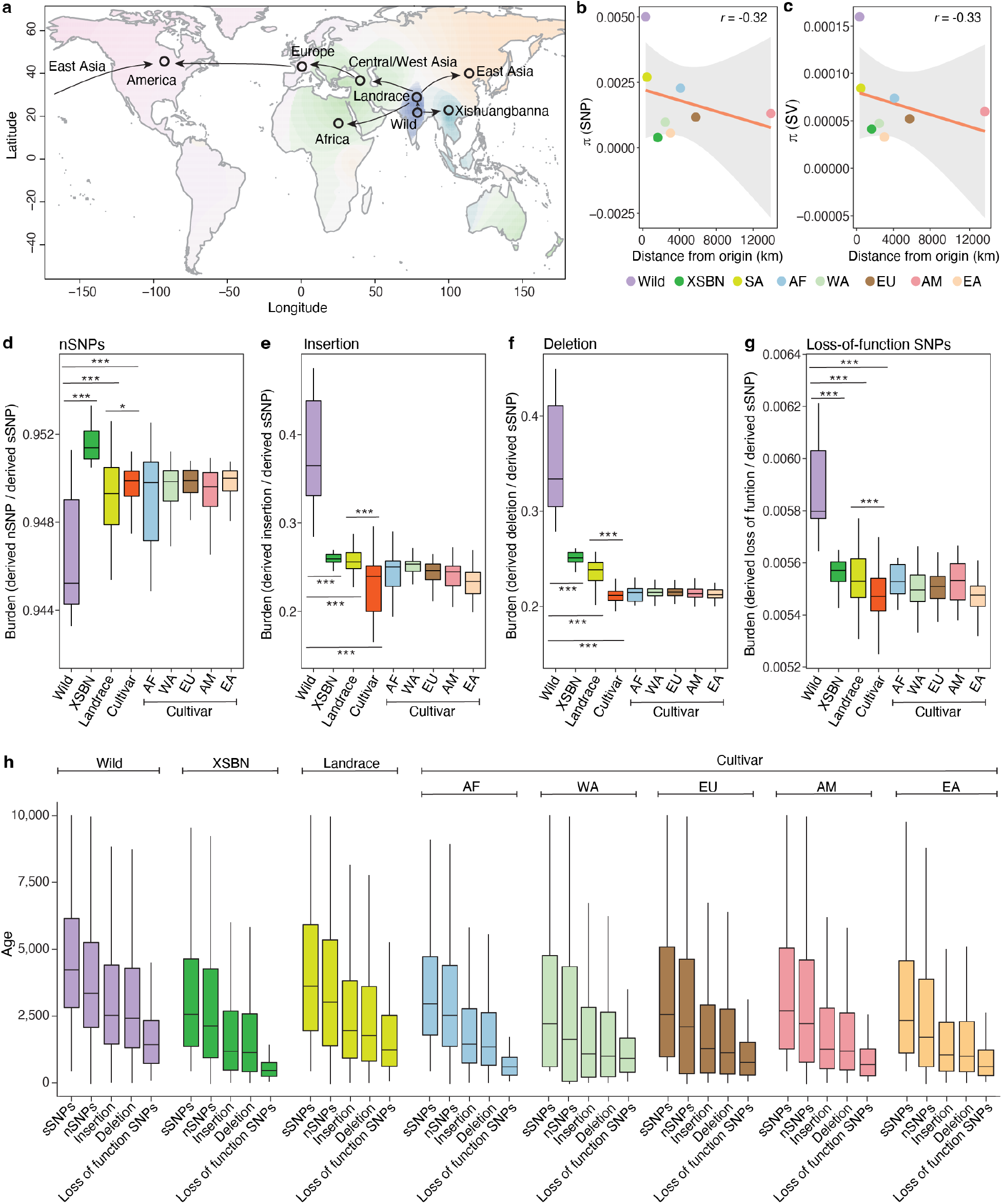
Spatiotemporal profiles of SV burden across different cucumber populations. **a**, Geographic representation of cucumber range expansion. Eight major geographical cucumber populations are shown on the map, with arrows indicating the putative expansion routes inferred from the population structure analysis. **b-c**, Correlation between nucleotide diversity (π) inferred from SNPs (**b**) and SVs (**c**) and the geographical distance of the eight cucumber populations from the center of origin. The red lines represent the linear regression trend and the shaded areas represent the confidence intervals of the linear regression. **d-g**, Burden of non-synonymous SNPs (nSNPs) (**d**), insertions (**e**), deletions (**f**), and loss-of-function SNPs (**g**) in different cucumber populations. *, ** and *** indicate significant differences at *P* < 0.05, 0.01, and 0.001, respectively. **h**, Age distribution of synonymous SNPs (sSNPs), nSNPs, insertions, deletions, and loss-of-function SNPs in different cucumber populations. For each boxplot, the lower and upper bounds indicate the first and third quartiles, respectively, the center line indicates the median, and the whiskers extend to 1.5× the interquartile range. XSBN, Xishuangbanna; AF, Africa; WA, Central/West Asia; EU, Europe; EA, East Asia; AM, America.

**Fig. 5.**
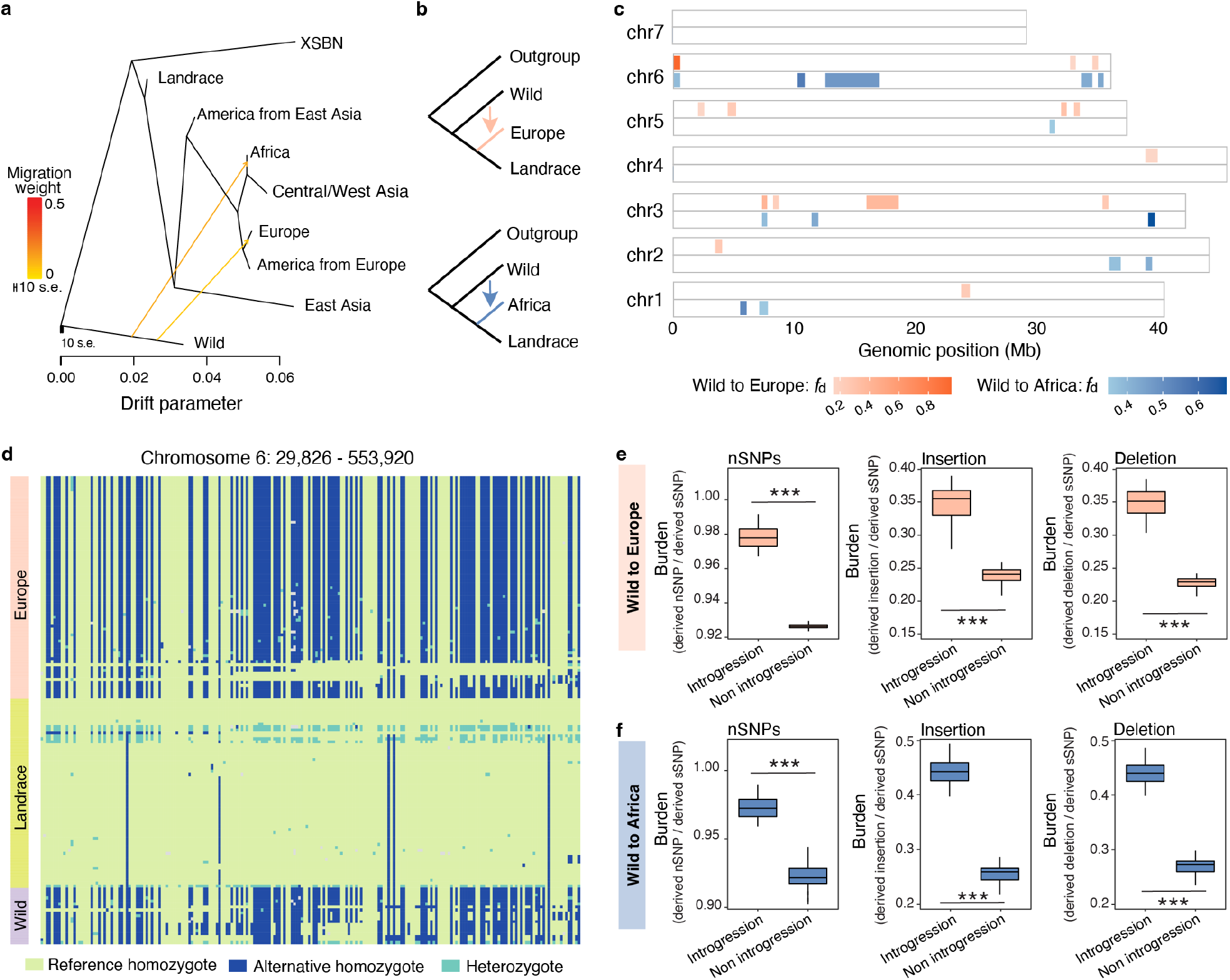
Adaptive introgressions in cucumber. **a**. Gene flow among different cucumber populations. Arrows indicate the direction and strength of gene flow. **b**, Phylogenetic topology illustrating introgression events from wild into European and African cucumbers. **c**, Genome-wide distribution of introgressed regions in European and African cucumbers. **d**, Heatmap of genotype profiles of an introgressed region from wild to the European population on chromosome 6. **e-f**, Comparative analysis of mutation burden in introgressed versus non-introgressed regions in European (**e**) and African (**f**) populations. For each boxplot, the lower and upper bounds indicate the first and third quartiles, respectively, the center line indicates the median, and the whiskers extend to 1.5× the interquartile range. *** indicates significant difference at *P* < 0.001.

Leveraging the advantages of graph-based pangenomes in representing SVs^5^, we used PanGenie^18^ to genotype SVs within the pangenome graphs for 414 accessions with high-depth genomic resequencing data. Among these accessions, 384 were newly sequenced in this study, with an average depth of 53.03×, including ten also sequenced for reference-level genome assemblies. The remaining 30 accessions were from a previous study^10^ (**Supplementary Table 6**). Together with the 39 genome assemblies, our data provided SV genotype information for a total of 443 accessions, including 18 wild accessions, 22 Xishuangbanna cucumbers, 66 landraces, and 337 cultivars. To validate the SV genotyping, we compared the SV genotypes generated from PanGenie with those identified from the pangenome graphs in 10 accessions that had both resequencing data and genome assemblies. The comparison demonstrated a high degree of concordance, with Pearson correlation coefficients of 0.97 for insertions and 0.96 for deletions, confirming the reliability of our SV genotyping (**Fig. 2e**).

We observed significant variation in both the number and length of SVs across different accessions and populations, with the wild populations exhibiting the highest number of SVs (**Fig. 2f, Supplementary Table 7** and **Supplementary Fig. 6**). Based on significantly different frequencies (FDR < 0.001 and fold change > 2) of SVs across breeding stages, we identified 7,517 SVs associated with domestication and 5,539 linked to the improvement process (**Supplementary Fig. 7**). These SVs affected 1,268 and 778 genes, respectively, including well-known genes involved in cucumber domestication and improvement, such as the fruit bitterness gene *bt*^19^ and the *CsYcf54* (*lgf*) gene, which influences fruit color^20^ (**Supplementary Fig. 7** and **Supplementary Tables 8, 9**).

**Fig. 6.**
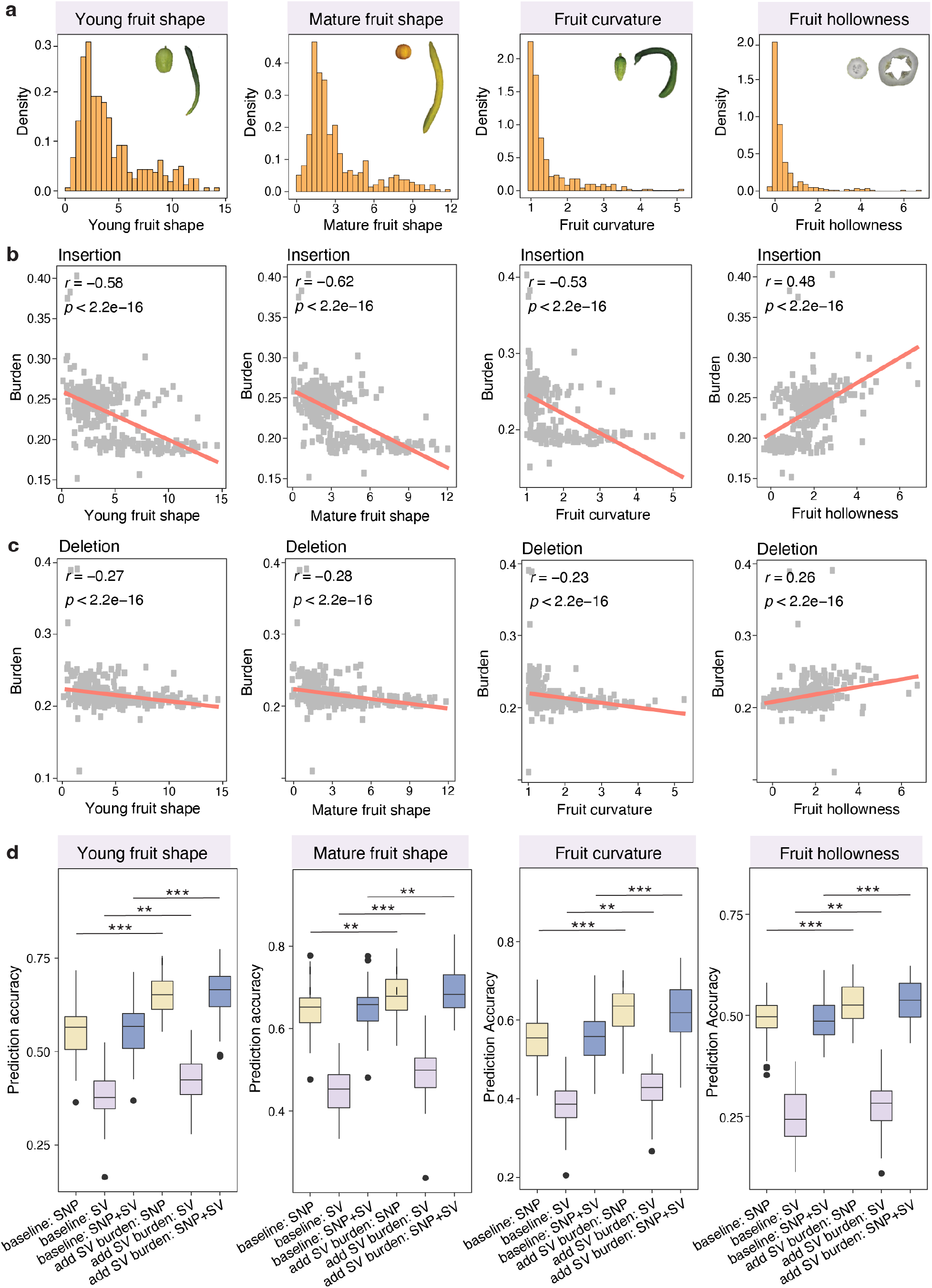
Genomic prediction incorporating SV burden information. **a**, Phenotypic distribution of young fruit shape, mature fruit shape, fruit curvature, and fruit hollowness. **b-c**, Correlations between insertion (**b**) and deletion (**c**) burden and the four fruit traits. **d**, Genomic prediction accuracies for the four traits using models with and without incorporating SV burden information. For each boxplot, the lower and upper bounds indicate the first and third quartiles, respectively, the center line indicates the median, and the whiskers extend to 1.5× the interquartile range. *, ** and *** indicate significant differences at *P* < 0.05, 0.01, and 0.001, respectively.

Phylogenetic analysis of cucumber accessions based on SVs clearly separated wild, landrace, and cultivated groups, as well as accessions from different geographic origins, consistent with the phylogenetic relationships inferred from SNPs (**Fig. 2j** and **Supplementary Fig. 8**), further supporting the reliability of SV genotyping. Population structure and principal component analyses using both SNPs and SVs revealed the spread of cucumbers from India to nearby regions, leading to the formation of the landrace group, with the landrace group subsequently diversifying globally (**Supplementary Figs. 9-11**). Interestingly, American cultivars clustered into two groups, one associated with East Asian cucumbers and the other with European cucumbers, consistent with recent findings that American cucumbers originated from Europe and were influenced by the intensive use of East Asian cucumbers in modern breeding^21,22^.

The fixation index (*F*_ST_) and nucleotide diversity (π), inferred from both SV and SNP data, underscored the distinct evolutionary trajectories of cucumbers across diverse environments (**Fig. 2h,i**). The wild species exhibited the highest nucleotide diversity, which decreased progressively through domestication and improvement, reflecting bottleneck events during these processes (**Fig. 2i**). Strong correlations between SVs and SNPs were observed for both *F*_ST_ (*r* = 0.920, *P* = 4.42.e-12) and π (*r* = 0.982, *P* = 1.47e-5), further confirming the reliability of SV genotyping and underscoring the role of SVs in capturing cucumber genomic diversity (**Fig. 2h,i**).

### SV dynamics during cucumber domestication and improvement

Our phylogenetic analysis traced the evolutionary transitions from wild cucumbers to landraces and, ultimately, to cultivated varieties. The Xishuangbanna population, however, represents a unique endemic lineage that does not contribute genetically to cultivated cucumbers (**Fig. 2g** and **3a**). Demographic history estimate revealed that landraces diverged from wild populations approximately 5,200 years ago (5,256.0 ± 507.3 years BP), marking the likely onset of cucumber domestication (**Fig. 3b,c**). This domestication process involved a prolonged but relatively mild bottleneck, followed by the spread of landraces across various regions, eventually leading to the development of cultivars through deliberate breeding efforts (**Fig. 3b,c**). We estimated that the split between cultivars and landraces occurred approximately 3,400 years ago (3,398.2 ± 626.1 years BP) (**Fig. 3b,c**). Unlike landraces, cultivars experienced a severe genetic bottleneck, reducing their effective population size to merely one-tenth of the original. However, with deliberate cultivation and adaptation to diverse environments, their effective population size recovered rapidly, enabling cultivars to emerge as the dominant local species (**Fig. 3b,c**).

We analyzed the unfolded site-frequency spectrum (SFS) during cucumber domestication and improvement for SVs (insertions and deletions), as well as for synonymous SNPs (sSNPs) and non-synonymous SNPs (nSNPs) (**Fig. 3d-f**). In both landrace and cultivar groups, most genetic variants were found at very low or very high frequencies, with fewer variants at intermediate frequencies, aligning with patterns observed in other crops such as rice^23^ and maize^24^ and consistent with the enhanced genetic drift associated with domestication and improvement bottlenecks. Notably, cultivars exhibited a higher proportion of fixed variants compared to landraces, suggesting that cultivars underwent a more severe genetic bottleneck, consistent with effective population size estimates (**Fig. 3b,d-f**). Additionally, SVs were skewed toward lower frequencies in the SFS compared to sSNPs in all three groups, indicating a lower proportion of fixed SVs relative to fixed sSNPs and nSNPs. The SFS of SVs differed significantly from that of sSNPs in all three groups (*P* < 0.05), suggesting that SVs may be predominantly deleterious or have higher mutation rates than SNPs, leading to a greater proportion of low-frequency variants (**Fig. 3d-f**). Furthermore, similar patterns between insertions and deletions were observed across different breeding states, suggesting comparable evolutionary dynamics and selective pressures acting on these two types of variants in cucumber (**Fig. 3d-f**). This contrasts with findings in grape which were based on short-read alignment to a single reference genome, where insertions exhibited a much lower proportion of fixed variants compared to deletions^25^. Given insertions and deletions are fundamentally similar in nature, they are expected to exhibit comparable evolutionary patterns^1^. This highlights the value of graph-based pangenomes, which substantially improve SV identification and enable more accurate biological insights.

To quantify the strength of selection acting on SVs during cucumber domestication, we estimated the distribution of fitness effects (DFE) based on population frequency data, using sSNPs as a neutral control^26^. This analysis encompassed a spectrum of selection intensities, ranging from negative to neutral and positive, with values of the strength of selection (*N*_e_*S*) near 0 indicating a neutral selection^27^. Our results revealed that SVs were subject to stronger purifying selection across wild, landrace, and cultivated populations compared to nSNPs, with a higher proportion of SVs under negative selection (*N*_e_*S* < 0) (**Fig. 3g-i**). For strongly deleterious mutations (*N*_e_*S* < -100), SVs experienced significantly greater selection intensity than nSNPs (*P* < 0.05), indicating their higher likelihood of being purged by natural selection due to their negative impact on plant fitness. In contrast, nSNPs showed a significant increase (*P* < 0.05) in neutral or near-neutral mutations (−1 < *N*_e_*S* < 1), suggesting a relaxation of selection pressure on mildly deleterious or neutral nSNPs, thereby facilitating the accumulation of these slightly harmful variants (**Fig. 3g-i**). This pattern indicates that while SVs are consistently purged by natural selection, slightly deleterious nSNPs may persist in domesticated cucumbers, potentially contributing to their genetic diversity.

Furthermore, nSNPs demonstrated higher levels of adaptive variation (α) compared to insertions and deletions, indicating that most adaptive changes in cucumbers are driven by SNPs rather than SVs (**Fig. 3j**). Across all variant types, the proportion of adaptive variation increased progressively from wild to landrace to cultivated populations, suggesting that domestication and improvement processes have selectively favored adaptive changes (**Fig. 3j**). In summary, the evolutionary dynamics of SVs during cucumber domestication indicate their strong purifying selection and their more constrained, often deleterious nature compared to SNPs. Domestication has primarily driven adaptive evolution through SNPs, with selection favoring beneficial SNPs while purging deleterious SVs, especially in cultivated cucumbers.

### SV dynamics during cucumber range expansion

Previous studies have shown that range expansion often intensifies genetic drift, leading to an increased frequency of deleterious mutations^28,29^. This accumulation reduces overall fitness and poses challenges to the persistence of newly established populations, a phenomenon known as expansion load^30^. To investigate the variation in SV load during cucumber range expansion, we analyzed eight populations along inferred migration routes based on phylogenetic and population structure analyses (**Fig. 2g, 3a** and **4a**). Our findings revealed a decline in genetic diversity, in terms of both SNPs and SVs, as cucumbers expanded from their origin in India (**Fig. 4b,c**). Populations at the expansion margins, particularly the Xishuangbanna population, exhibited the lowest genetic diversity (**Fig. 4b,c**). This reduction in genetic diversity along the expansion route suggests that cucumbers may have experienced expansion load.

Next, we quantified mutation load by calculating the ratio of derived nonsynonymous or SV alleles to derived synonymous alleles^31^. Our analysis revealed that the mutation load of nSNPs, which are often mildly deleterious, increased significantly during domestication and population expansion (**Fig. 4d**). Compared to the wild population, landraces exhibited a 0.44% increase in mutation load (*P* = 2.59e-4), supporting the ‘cost of domestication’ hypothesis^23^. Additionally, cultivars showed a significantly higher mutation load than landraces (*P* = 0.0196). While no significant differences were detected among the five cultivated groups, the Xishuangbanna population, a marginal group, exhibited the highest mutation load (**Fig. 4d**). This finding aligns with the expansion load hypothesis, highlighting the role of genetic drift in accumulating slightly deleterious or nearly neutral nSNPs. Furthermore, the observed decreases in genetic diversity (**Fig. 4b**) during domestication and expansion suggest a reduction in effective population size, as these two factors are typically positively correlated^32^. A reduced effective population size weakens purifying selection, allowing slightly deleterious mutations to persist and accumulate through genetic drift^26,33^. Combined with historical bottlenecks during domestication, these factors likely contributed to the accumulation of SNP expansion load.

In contrast to nSNPs, we observed a reduction in SV load during population expansion (**Fig. 4e**,**f**). Compared to the wild population, SV load in landraces and cultivars decreased by 29.86% (*P* = 4.37e-8) and 34.45% (*P* = 3.00e-9) for insertions, and by 28.51% (*P* = 5.37e-8) and 36.41% (*P* = 4.58e-9) for deletions, respectively (**Fig. 4e,f**). Deleterious SVs, similar to loss-of-function SNPs (defined as those that introduce or remove stop codons), showed no signs of expansion load, suggesting that high-impact SVs are consistently purged by purifying selection (**Fig. 4g**).

We further analyzed the age distribution of SNPs and SVs, revealing that sSNPs and nSNPs exhibited a relatively older age distribution (**Fig. 4h**). This suggests that these SNPs have accumulated over the course of cucumber evolution under relatively weaker purifying selection. In contrast, insertions and deletions displayed a younger age distribution (**Fig. 4h**), especially in landrace and cultivar populations. This indicates that SVs are more recent and likely subject to stronger purifying selection during domestication and geographic expansion. This observation is consistent with the reduced SV burden observed in cultivars, suggesting that recently emerged deleterious SVs are more effectively purged during the expansion process, thereby minimizing their accumulation in cultivated varieties. Additionally, loss-of-function SNPs exhibited the youngest age among all variants (**Fig. 4h**), highlighting their highly deleterious nature. These findings underscore the critical role of purifying selection in reducing the burden of SVs and loss-of-function SNPs, shaping the genetic landscape by purging highly deleterious variants.

### SV dynamics in cucumber introgression

Introgression, the transfer of alleles from a donor to a recipient lineage, is a pervasive evolutionary process with significant implications for population fitness and genomic architecture^34^. While maladaptive alleles introduced via introgression are often purged by natural selection, adaptive variants can enhance local adaptation and reduce genetic load^35^. Understanding the genomic patterns and consequences of introgression is important for improving crop traits such as stress tolerance, which is crucial for food security^36^. Using Treemix^37^, we detected significant gene flow from wild cucumber to the African and European cucumbers (**Fig. 5a**). Further analysis with ABBA-BABA statistics^38^ identified 14 introgressed genomic regions containing 1,043 genes in the European population and 13 regions encompassing 1,165 genes in the African population (**Fig. 5b,c** and **Supplementary Table 10**). Notably, an introgressed region on chromosome 6 in the European group, exhibiting the strongest signal (**Fig. 5c**,**d**), harbored a CBL-interacting protein kinase gene, whose homologs are key regulators of plant responses to environmental stress and growth development^39^. In addition, a prominent introgressed region in the African population on chromosome 3 contained a homolog of the wheat disease resistance gene *Lr10* (ref. ^40^) (**Supplementary Fig. 12**). This introgression landscape highlights the importance of leveraging genetic diversity from wild populations to enhance cucumber breeding and improvement efforts. To investigate the genetic consequences of introgression in cucumber, we compared the SFS of SNPs and SVs in introgressed versus non-introgressed regions in the European population, using sSNPs as the neutral reference. Interestingly, introgressed regions exhibited lower frequencies in the SFS compared to non-introgressed regions, indicating a lower proportion of fixed alleles in introgressed regions (**Supplementary Fig. 13**). Moreover, introgression appeared to increase the deleterious load, as evidenced by a significantly higher ratio of derived alleles in nSNPs and SVs relative to sSNPs in introgressed regions compared to non-introgressed regions (*P*< 2.2e-16) (**Fig. 5e**). This suggests that introgressed regions harbor a higher number of deleterious variants, likely due to linkage drag that introduces harmful variants alongside beneficial traits. Specifically, numerous low-frequency deleterious variants in wild populations may become largely fixed after introgressed into cultivars, substantially elevating the deleterious load in introgression regions. Similar patterns were observed for introgressions in the African population, further supporting these conclusions (**Fig. 5f** and **Supplementary Fig. 13**).

### Incorporating SV burden improves genomic prediction

Genomic prediction has emerged as an invaluable tool in plant and animal breeding for estimating the genetic potential of individuals^41^. Incorporating deleterious mutations into genomic prediction models has been shown to improve predictive accuracy in crops such as maize^42,43^ and potato^44^. However, the impact of SVs on genomic prediction accuracy in cucumber remained unexplored. To address this gap, we analyzed 21 key fruit quality and yield traits in a core cucumber collection of 374 accessions^45^. We found that four traits were significantly correlated with SV burden (*P* < 2.2e-16) (**Fig. 6a-c** and **Supplementary Fig. 14**). Specifically, traits such as young fruit shape, mature fruit shape, and fruit curvature exhibited a significant negative correlation with SV burden, while fruit hollowness showed a significant positive correlation (**Fig. 6a-c**).

Given the significant correlation between mutation burden and agronomic traits, we conducted genomic prediction for these four key traits (**Fig. 6d**). Using a baseline model that accounts for genomic relationships without considering deleterious mutations, the prediction accuracies for young fruit shape, mature fruit shape, fruit curvature, and fruit hollowness using SNPs alone were 0.550, 0.644, 0.551, and 0.490, respectively (**Fig. 6d** and **Supplementary Table 11**). When using the baseline model with SVs alone, prediction accuracies were substantially lower than those achieved with SNPs alone. Furthermore, combining SNPs and SVs in the baseline model did not substantially improve prediction accuracies compared to using SNPs alone (**Supplementary Fig. 15**). This may be attributed to the fact that SNPs and SVs are often linked, and incorporation of SVs may not provide substantial additional information beyond what is already captured by genome-wide high-density SNPs.

Furthermore, adding SNP burden to the baseline model did not improve prediction accuracy compared to the baseline model without SNP burden (**Supplementary Fig. 16** and **Supplementary Table 11**). In contrast, incorporating SV burden into the prediction model enhanced prediction accuracy for the four traits significantly correlated with SV burden. Specifically, prediction accuracy increased by 19.36% for young fruit shape, 4.51% for mature fruit shape, 13.57% for fruit curvature, and 8.22% for fruit hollowness. Moreover, adding SV burden to models using SVs or combined SNPs and SVs improved prediction accuracy compared to the baseline models using only SVs or combined SNPs and SVs (**Fig. 6d** and **Supplementary Table 11**). Interestingly, combining SNP and SV burden did not yield further improvements over SV burden alone, suggesting that SNP burden may have limited influence on these traits, potentially diluting the impact of SV burden. This was further supported by genomic predictions on 17 additional traits not correlated with SV burden, where no significant improvements in prediction accuracy were observed (**Supplementary Fig. 16**).

## Discussion

SVs represent a relatively unexplored aspect of plant genome variation, with limited knowledge regarding their population dynamics^3^. In this study, we utilized a graph-based pangenome approach to comprehensively and accurately identify 171,892 SVs across 39 reference-level genome assemblies in cucumber. By genotyping these SVs in 443 accessions spanning wild, landrace, and cultivated cucumbers, we developed a robust resource for investigating SV dynamics and their contributions to cucumber evolution and breeding. The newly developed reference-level genome assemblies and the comprehensive population-level SV dataset provide unprecedented resources for future comparative and functional genomic studies, as well as for advancing cucumber improvement efforts.

Our findings illuminate the complex dynamics of SVs, emphasizing their significant roles in domestication, geographic expansion, and introgression. During domestication, adaptive evolution was predominantly driven by SNPs, with selection favoring beneficial SNPs while purging deleterious SVs, reflecting the stronger deleterious effects often associated with SVs. During cucumber range expansion, SVs exhibited an intermediate deleterious effect, falling between nSNPs and loss-of-function SNPs. Recently emerged deleterious SVs were more effectively purged during expansion, minimizing their accumulation in cultivated varieties. In contrast, nSNPs showed clear signs of expansion load in marginal populations, likely due to genetic drift. During introgression, particularly from wild cucumbers into African and European populations, the introduction of beneficial alleles was accompanied by a burden of linked deleterious mutations through hitchhiking. This led to an increased deleterious mutation load, which is in contrast to findings in maize and bread wheat, where introgressions from wild teosinte and wild emmer, respectively, reduced the deleterious allele burden^46,47^. This discrepancy may reflect differences in the timing and extent of recombination following introgression. In maize and bread wheat, introgressions occurred approximately 6,000–9,000 years ago^48,49^, allowing sufficient time for recombination to unlink beneficial alleles from deleterious ones. By contrast, cucumber cultivars originated around 3,500 years ago, suggesting that introgressions occurred more recently, leaving insufficient time for recombination to purge deleterious variants. Therefore, these introgression-associated deleterious variants represent potential targets for genomic design of cucumber breeding, particularly in efforts to reduce the burden of harmful mutations.

Incorporating SV burden into genomic prediction models significantly improved prediction accuracy for several key agronomic traits, highlighting the unique and complementary information provided by SVs. However, our findings also revealed that combining SNPs and SVs in baseline models did not always yield substantial improvements, likely due to linkage between these two variation types. The integration of SV burden is particularly valuable for traits strongly influenced by SVs, enabling breeders to prioritize deleterious variants that impact phenotypes while maintaining breeding efficiency. These findings underscore the need for trait-specific genomic strategies that align prediction models with the genetic architecture of target traits, reducing redundancy and optimizing breeding outcomes.

## Methods

### Genome sequencing, assembly and annotation

Cucumber plants were grown in the greenhouse at Boyce Thompson Institute in Ithaca, New York, with a 16-h light period at 20 °C (night) to 25 °C (day). High-molecular-weight genomic DNA was extracted from fresh young leaves of cucumber plants. HiFi SMRTbell libraries were constructed following the standard protocol provided by PacBio and sequenced on a PacBio Sequel II platform using the circular consensus sequencing (CCS) mode to generate HiFi reads, which were *de novo* assembled into contigs using hifiasm^50^ (v0.19.6-r595) with default parameters. The assembled contigs were anchored and oriented into seven cucumber chromosomes using RagTag^51^ (v.2.0.1), with the Chinese Long 9930 genome serving as the reference. Large structural variations such as inversions or translocations were manually inspected and corrected by checking the HiFi read alignments.

Repeat sequences across the 39 cucumber genome assemblies were identified using RepeatModeler^52^ (v.2.0.4) and RepeatMasker^53^ (v.4.1.1). Protein-coding genes were initially predicted from the repeat-masked genomes using the MAKER pipeline^54^ (v.3.01.03), which integrates *ab initio* predictions, transcriptome evidence, and homologous protein evidence. Specifically, AUGUSTUS^55^ (v.3.4.0) and SNAP^56^ (v.2006-07-28) were used for *ab initio* gene predictions. RNA-Seq data from diverse cucumber tissues, including root, stem, leaf, flower, and fruit^57^, were used for transcriptome evidence. To provide protein homology evidence, protein sequences from Arabidopsis (TAIR10), *Cucumis melo*^58^, *C. hystrix*^59^, *C. hardwickii*^10^, cucumber^12^ (9930), cucumber (Gy14 v2) (http://cucurbitgenomics.org/v2/), and the UniProt (Swiss-Prot plant division) database were aligned to the genomes using Spaln^60^. Subsequently, Liftoff^61^ (v.1.5.1) was employed to transfer predicted gene models across assemblies. Furthermore, Helixer^62^ (v.0.3.2), a deep learning-based framework for cross-species gene annotation, was also applied. Finally, the results from all approaches were integrated using EVidenceModeler^63^ (v.1.1.1) to generate the final set of predicted protein-coding genes for each genome assembly.

### Gene-based pangenome construction and phylogenetic analysis

Protein sequences from the 39 cucumber genomes were clustered into gene families (pan-gene clusters) using OrthoFinder^64^ (v.2.5.5). Based on their presence across the accessions, pan-gene clusters were categorized into four distinct groups: core (present in all accessions), soft core (present in more than 90% but not all accessions; 35 ≤ n < 39), dispensable (present in more than one but less than 90% of accessions; 2 ≤ n < 35), and private (present in only one accession). Nucleotide diversity (π) for each gene in the pangenome was calculated based on multiple sequence alignments using MAFFT^65^ (v.7.526), according to the formula: 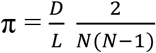, where D represents the number of variable sites, L is the length of the conserved alignment, and N is the number of sequences. The ratio of nonsynonymous to synonymous substitutions (*K*a/*K*s) for each gene in the pangenome was calculated using KaKs_Calculator^66^ (v. 2.0).

A total of 12,655 single-copy orthologous genes across the 39 genomes were used to construct the phylogenetic tree through a multispecies coalescent analysis in BEAST2 (ref. ^67^) with default parameters. The resulting tree was visualized using ggtree^68^.

### Graph-based pangenome construction

The cucumber graph-based pangenome was constructed using the Minigraph-Cactus package^69^ with the 39 reference-level genome assemblies. The construction process began with the AM716 genome as the initial graph, to which the remaining 38 assemblies were sequentially mapped. Graphs for individual chromosomes were then merged into a comprehensive whole-genome graph, which was indexed and exported to VCF format using the vg toolkit^70^. SVs, including insertions and deletions, were defined relative to the AM716 reference genome. The final graph was filtered using vcfbub (https://github.com/pangenome/vcfbub) with parameters ‘-l 0 -r 10000’.

### Short-read sequencing, SNP calling, and SV genotyping

Genomic DNA was isolated from young fresh leaf tissues using the Omega Mag-Bind Plant DNA DS Kit (M1130, Omega Bio-Tek, Norcross, GA), following the manufacturer’s instructions. Shotgun libraries were constructed from the extracted DNA and sequenced on the BGISEQ-500 platform to generate 150-bp paired-end reads. Raw sequencing reads were processed to remove adaptor and low-quality sequences using Trimmomatic^71^ (v.0.39). For SNP calling, the cleaned reads were aligned to the AM716 genome using BWA-MEM^72^ (v.0.7.17-r1188) with default parameters. Duplicate read pairs were marked, and variants were called using the Sentieon package (https://www.sentieon.com/). SNPs were filtered using GATK^73^ with parameters ‘QD < 2.0 || FS > 60.0 || MQ < 40.0 || MQRankSum < -12.5 || ReadPosRankSum < -8.0’. Additional filtering steps retained only biallelic SNPs with minor alleles present in at least two accessions and supported by a read depth between 0.67× and 1.5× of the genome-wide mode depth. SVs in the graph-based pangenome were genotyped in the resequenced cucumber accessions using PanGenie^18^ (v.3.0.2) with short sequencing reads, employing default parameters. SNPs and SVs were annotated using SnpEff^74^ (v.5.1).

### Population genomic analyses

Maximum-likelihood phylogenetic trees were constructed separately with the complete set of SVs and 294,365 SNPs located at fourfold degenerate sites (4DTv), using RAxML^75^ under the GTRGAMMA model with 100 bootstrap replicates. Three melon accessions (GenBank accession numbers: SRR8925010, SRR8925011, and SRR8925012) were used as the outgroup. The resulting phylogenetic trees were visualized using iTOL^76^. Population structure was inferred using ADMIXTURE^77^ with SNPs having a minor allele frequency (MAF) ≥ 0.05 and linkage disequilibrium (LD) ≤ 0.2. Geographical projections of the population structure were generated using the HCLUSTER package and visualized with the TESS3 package^78^. Principal component analysis (PCA) was performed using PLINK^79^. Nucleotide diversity (π) within each population and fixation index (*F*_ST_) between populations were calculated with SNPs and SVs, separately, using VCFtools^80^ (v.0.1.15) in 100-kb sliding windows with a 10-kb step size.

To identify SVs under selection during domestication and improvement, the occurrence frequencies of SV alleles were calculated across three groups: wild, landrace, and cultivar. The significance of SV allele frequency differences between wild and landrace (for domestication) and between landrace and cultivar (for improvement) was assessed using Fisher’s exact test. The resulting raw p-values were corrected for multiple testing using a false discovery rate (FDR) approach. SVs with FDR < 0.001 and fold change > 2 in allele frequencies were identified as being under selection.

### Reconstruction of demographic history and estimation of variant age

Effective population sizes were inferred using SMC++^81^ (v.1.15.4.dev18+gca077da). Due to the inbred nature of cucumber lines, the genome of each accession was treated as a single haplotype, with occasional heterozygous sites resolved by randomly selecting one allele. For each population, haplotypes from different accessions were combined to create pseudo-diploid genotypes. A total of 15 pseudo-diploid genotypes were randomly selected for the SMC++ analysis, using a mutation rate of 6.5×10^−9^ (ref. ^82^) and a generation time of 1 year. The SMC++ analysis was repeated 20 times for each population, with pseudo-diploid genotypes resampled in each iteration.

The ages of SNPs and SVs were estimated using GEVA^83^ (v.1beta), applying the same mutation rate of 6.5×10^−9^ (ref. ^82^). SNP and SV ages were computed using the joint clock model, which integrates both mutation and recombination clock models to provide more accurate estimations of variant ages.

### Site-frequency spectrum and distribution of fitness effects

To derive the unfolded SFS of SNPs and SVs, the ancestral and derived alleles for each variant were determined using *Cucumis melo* (melon)^58^ and *C. hystrix*^59^, two closely related species of cucumber, as outgroup species. SNP and SV alleles shared among cucumber, melon, and *C. hystrix* were identified as ancestral alleles. For each variant site, the derived allele was identified as the alternative allele in cucumber that differed from the ancestral allele. All SVs were identified relative to the AM716 genome, which served as the coordinate reference during graph construction. Accordingly, SV load per accession was calculated as the number of derived SVs mapped with respect to AM716 coordinates.

The distribution of fitness effects (DFE) and the proportion of adaptive variants (α) for nSNPs and SVs were estimated using the polyDFE program^84^ (v.2.15) based on the unfolded SFS. The 95% confidence intervals of the α estimates were derived from discretized DFEs inferred through 20 bootstrap replicates.

### Detection of gene flow

Gene flow between the eight cucumber groups was detected using TREEMIX^37^ (v.1.13). The optimal number of migration edges (m = 2) was obtained using the linear modeling estimate in the OptM R package (v0.1.6; https://cran.r-project.org/web/packages/OptM) with the number of migrations set from 0 to 10 and each test with 20 interactions. Genome-wide introgressions were detected by calculating the *f*_d_ statistic^38^ in sliding windows with 1,000 SNPs and a minimum of three valid sites per window. The *f*_d_ statistic was calculated using the landrace group as P1, the European or African population as P2, and the wild population as P3.

### Genomic prediction

Genomic prediction was conducted on a total of 21 agronomic traits using the R package rrBLUP^85^, incorporating SNP and SV burden information. Phenotypic data for these traits in the cucumber core collection were obtained from our recent study^45^. The prediction accuracy of rrBLUP was evaluated using five-fold cross-validation, repeated 50 times to ensure robustness. The prediction accuracy was calculated by averaging the Pearson correlation coefficients (*r*) between predicted and observed phenotypic values across the cross-validation replicates. SNPs were filtered to exclude closely linked sites using LDAK^86^ (v.5.2). Various genomic prediction models were tested within the rrBLUP framework, including those using only SNPs (baseline: SNP), only SVs (baseline: SV), a combination of SNPs and SVs (baseline: SNP+SV), and models that incorporated additional SNP, SV, and SNP+SV burden information.

## Data availability

Raw genome resequencing reads have been deposited in the NCBI BioProject database under the accession number PRJNA1192329. Raw HiFi reads and genome assemblies have been deposited in the NCBI Bioproject database under the accession number PRJNA844366. Genome assemblies and annotations, SNPs, small indels and SVs in VCF format are available at CuGenDBv2 (http://cucurbitgenomics.org/v2/ftp/pan-genome/cucumber/).

## Author contributions

Z.F. and Y.X. conceived the project. Z.F. designed and supervised the study. X.Z., J.Y., H.S. and S.W. contributed to genome assembly and annotation, pangenome construction, and SV genotyping. X.Z. performed population genetic analyses. X.Z., J.Zhao and Y.Z. contributed to genomic prediction analysis. R.G., S.A.H. and Y-C.L. contributed to sample collection, DNA extraction and phenotyping. R.T.D. and F.C. helped develop the population for sequencing. J.Zhang, Y.X., Y.W. and Z.F. coordinated genome sequencing. X.Z. wrote the paper. Z.F., Z.Z, S.H., Y.W., R.G. and Y.X. revised the paper.

## Acknowledgements

The authors thank Savannah Beyer (USDA-ARS) for technical help in developing the core collection. This research was supported by grants from USDA National Institute of Food and Agriculture Specialty Crop Research Initiative (2015-51181-24285 and 2020-51181-32139).

