## Supplementary figures for "Graph-based pangenome reveals structural variation dynamics during cucumber breeding"

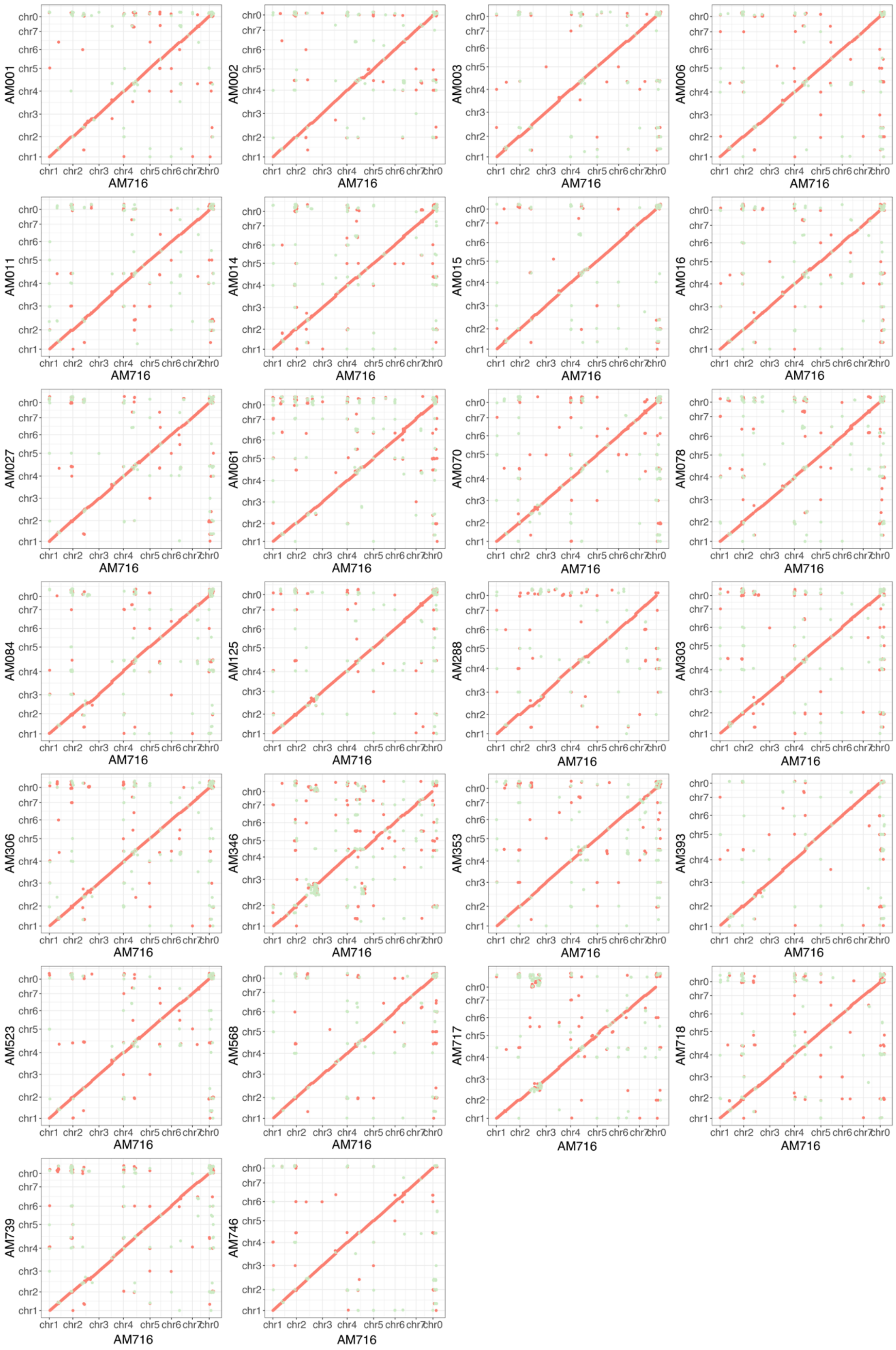


**Supplementary Fig. 1. Collinearity between the AM716 genome and the other 26 cucumber genomes assembled from PacBio HiFi reads.**

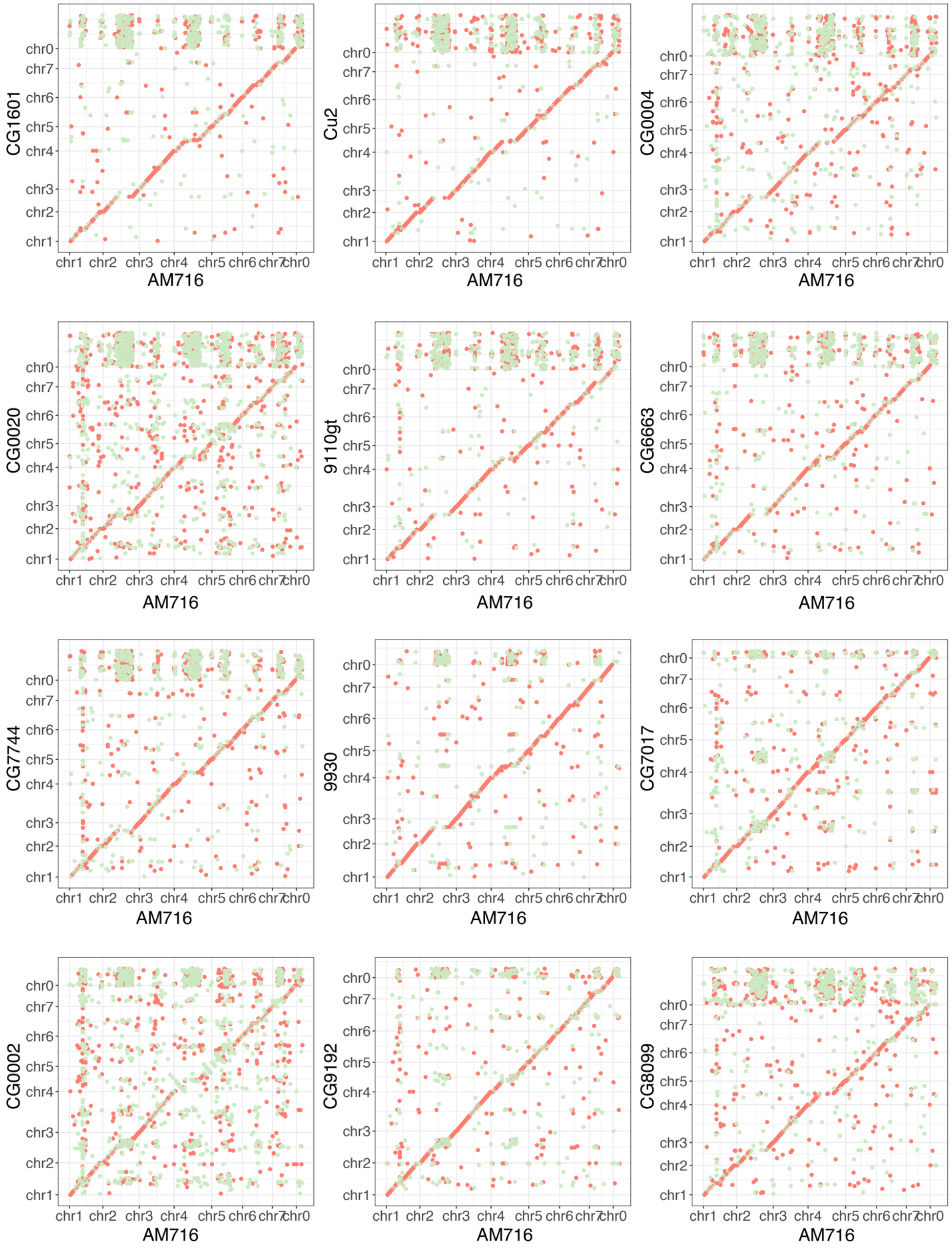


**Supplementary Fig. 2. Collinearity between the AM716 genome and 12 previously reported chromosome-scale cucumber genome assemblies.**

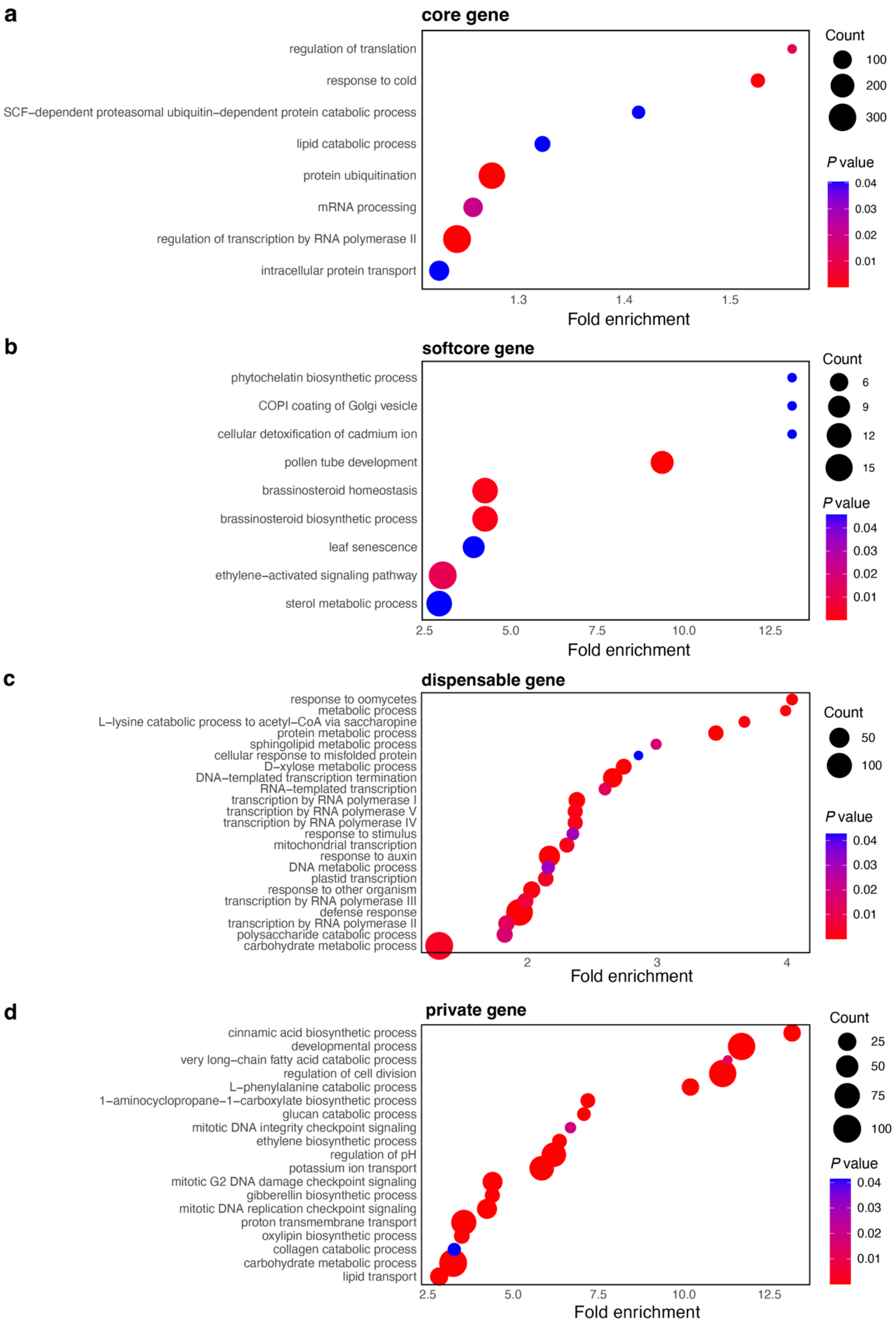


**Supplementary Fig. 3. Functional enrichments of core (a), softcore (b), dispensable (c), and private (d) genes in the cucumber gene-based pangenome.**

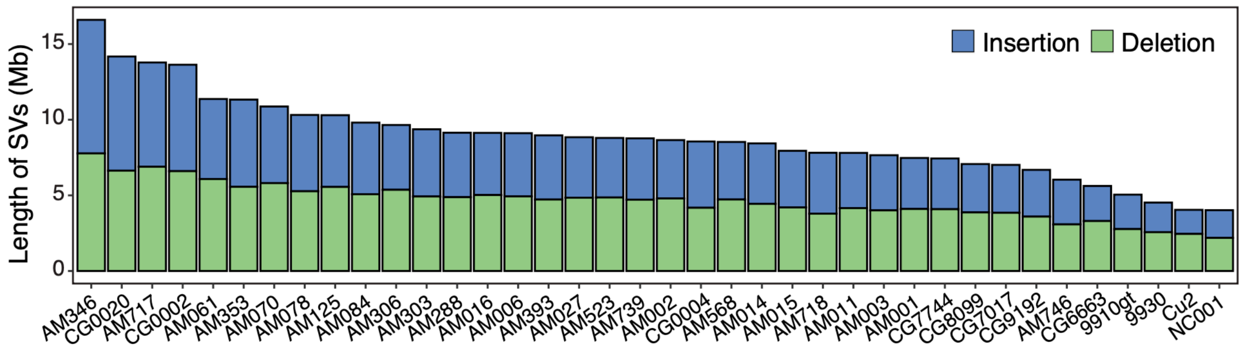


**Supplementary Fig. 4. Distribution of SVs across cucumber accessions.** Total length of all SVs in each accession.

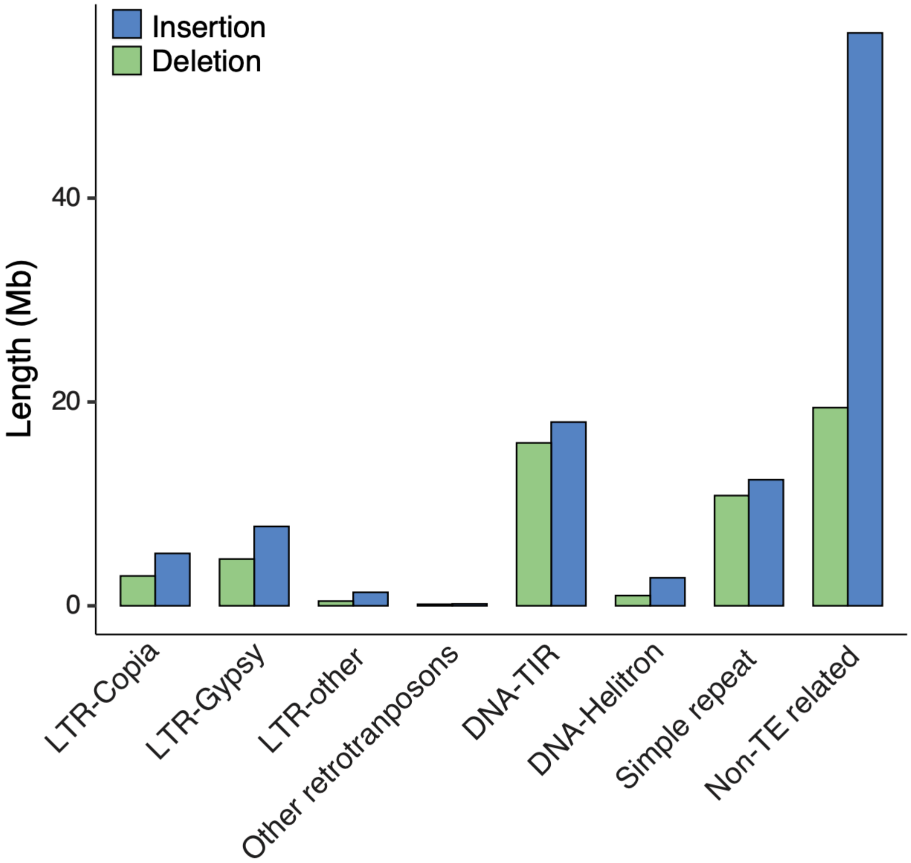


**Supplementary Fig. 5. Length distribution of SVs in cucumber derived from different types of repetitive sequences.**

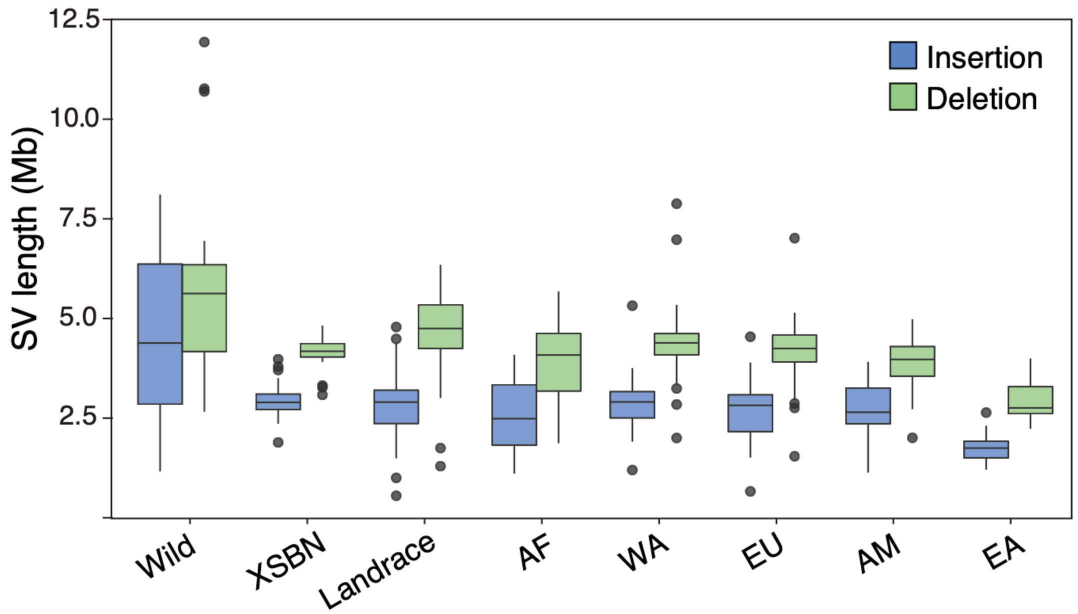


**Supplementary Fig. 6. Total length of SVs across different cucumber populations.** For each boxplot, the lower and upper bounds indicate the first and third quartiles, respectively, the center line indicates the median, and the whiskers extend to 1.5× the interquartile range. XSBN, Xishuangbanna; AF, Africa; WA, Central/West Asia; EU, Europe; EA, East Asia; AM, America.

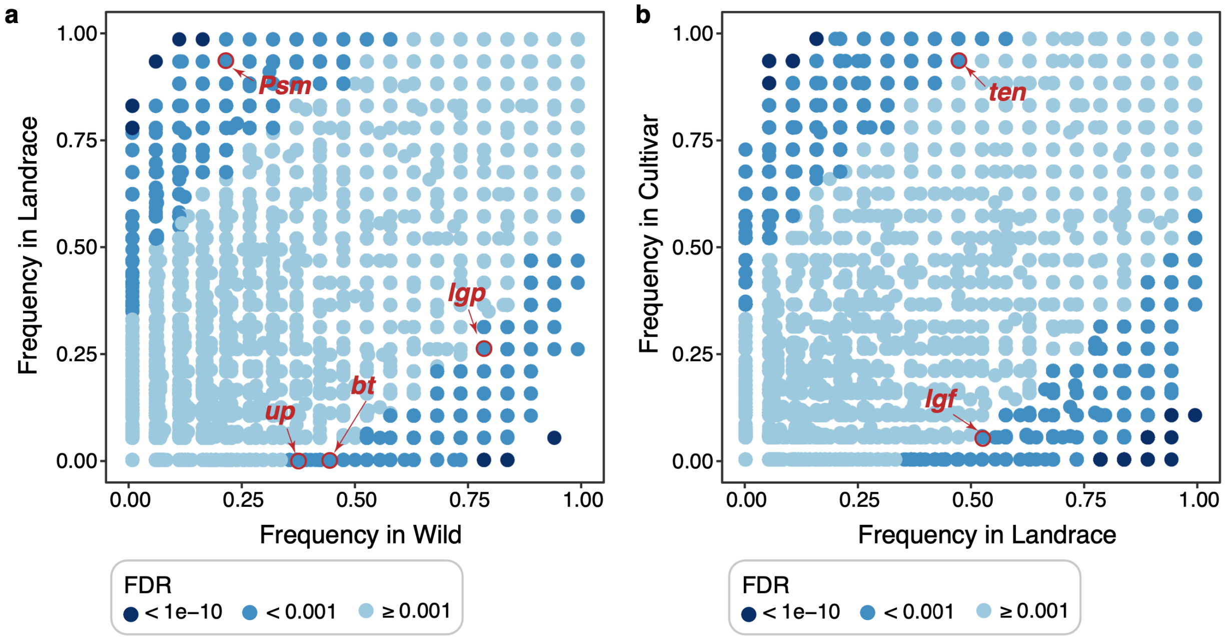


**Supplementary Fig. 7. SVs under selection during cucumber domestication and improvement. a,** Comparison of SV occurrence frequencies between wild and landrace populations (domestication). **b,** Comparison of SV occurrence frequencies between landrace and cultivar populations (improvement). SVs associated with known genes regulating key agronomic traits, including *Psm* (*Paternal sorting of mitochondria*), *lgp* (*light green peel*), *bt* (*bitter fruit*), *up* (*upward-pedicel*), *ten* (*tendril-less*), and *lgf* (*light green fruit*), are shown.

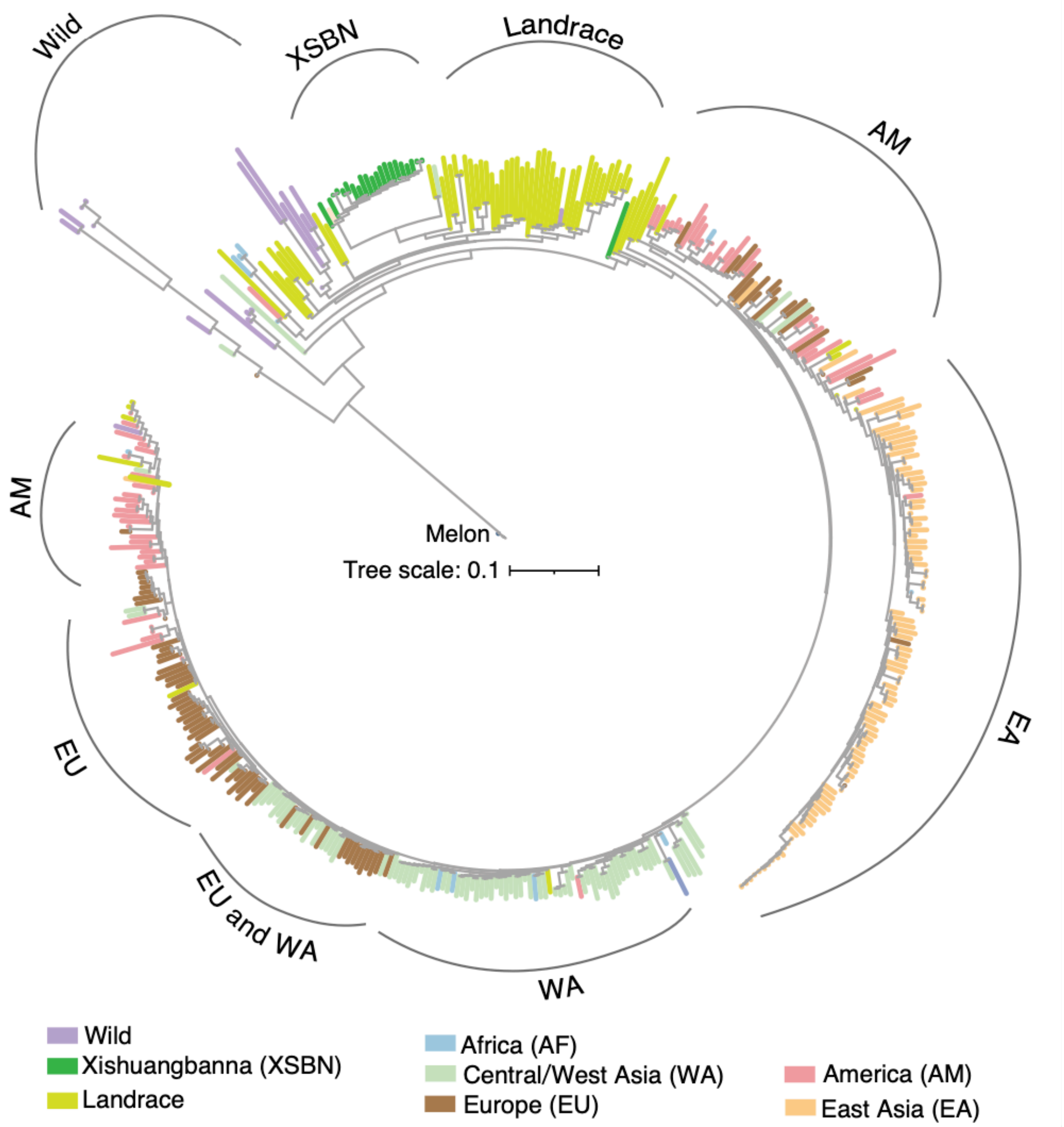


**Supplementary Fig. 8. Phylogenetic relationships of cucumber accessions based on SNPs.** The curved arcs on the periphery indicate clade-level groupings, each labeled by the predominant population identity of its member accessions.

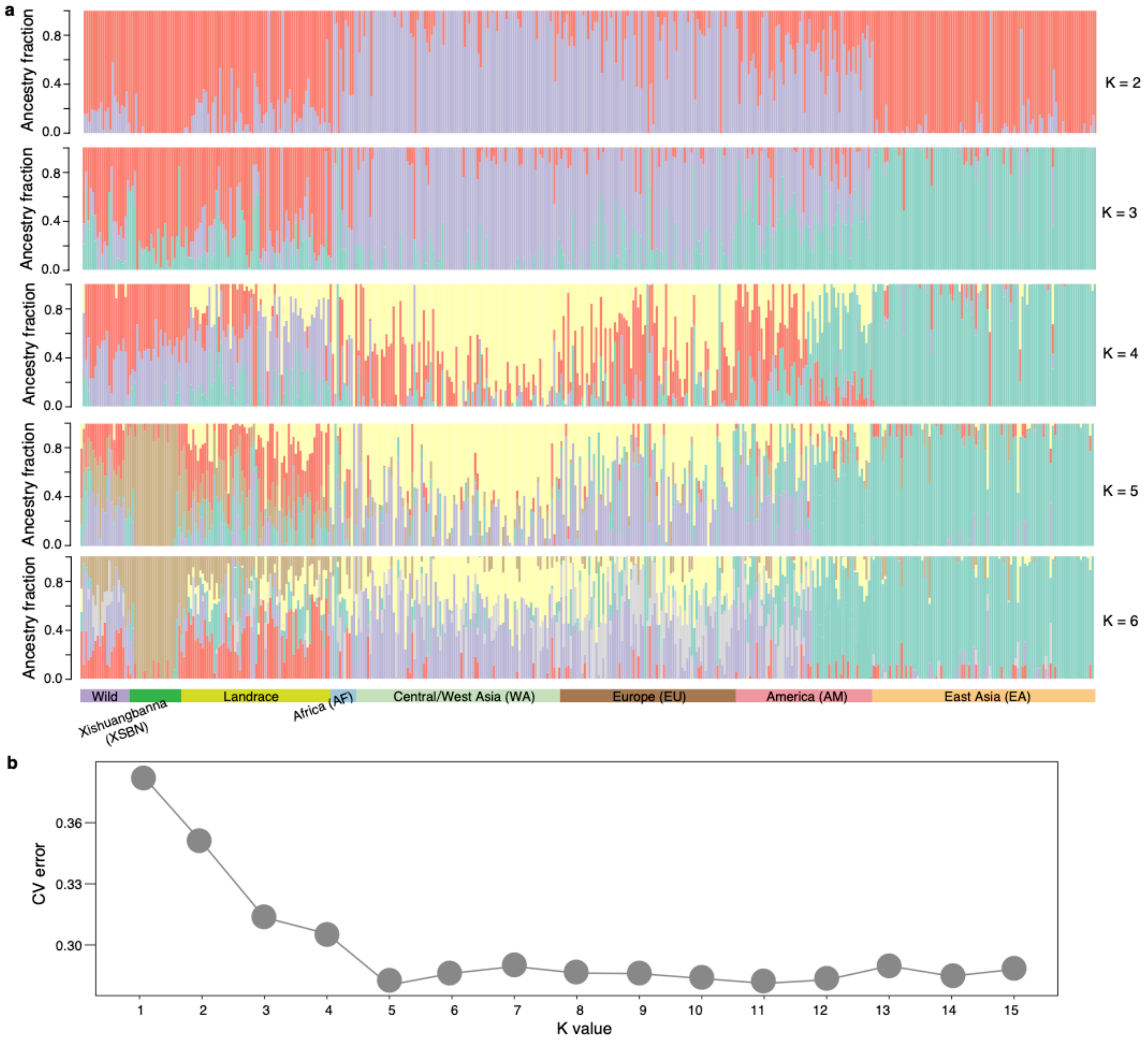


**Supplementary Fig. 9. Population structure of cucumber accessions based on SNPs. a,** Ancestry coefficient analysis with $K = 2$ to 6. **b,** Cross-validation (CV) error plotted against the number of clusters ($K$) used to infer population structure in cucumber.

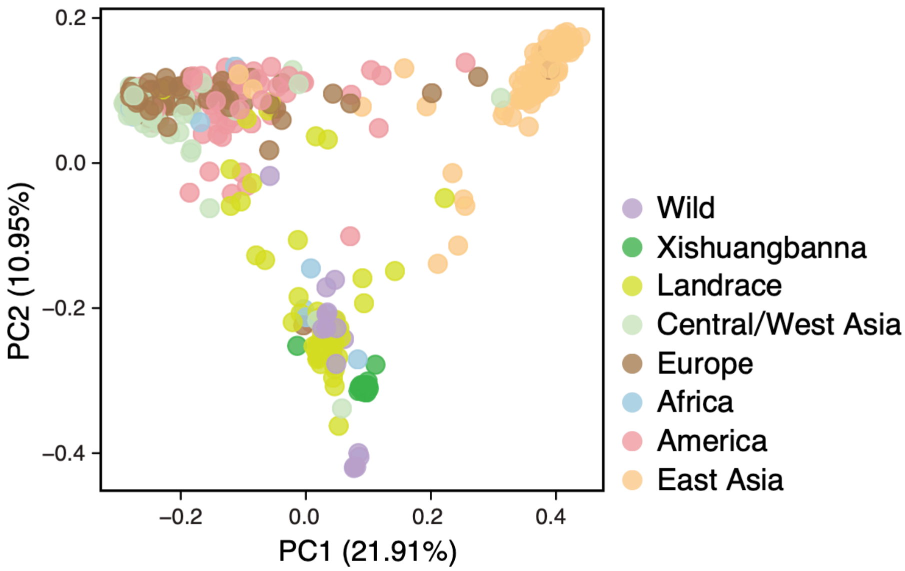


**Supplementary Fig. 10. Principal component analysis (PCA) of cucumber accessions based on SVs.**

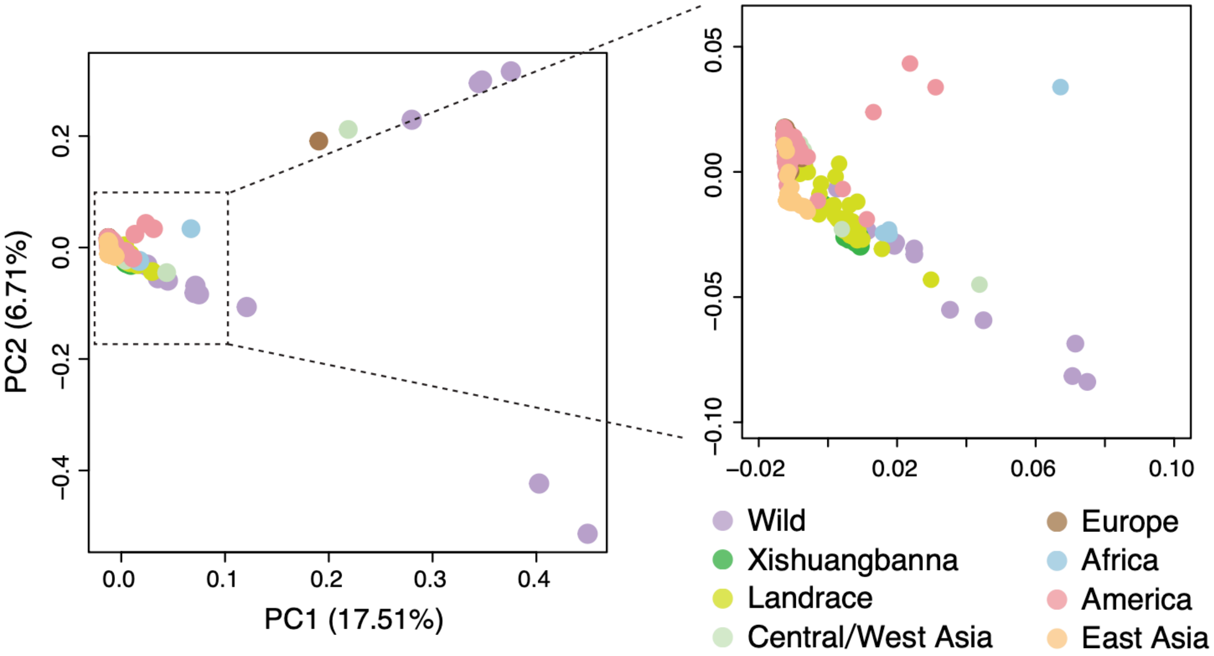


**Supplementary Fig. 11. Principal component analysis (PCA) of cucumber accessions based on SNPs.** The right panel provides a magnified view of the cluster indicated by the dotted box in the left panel.

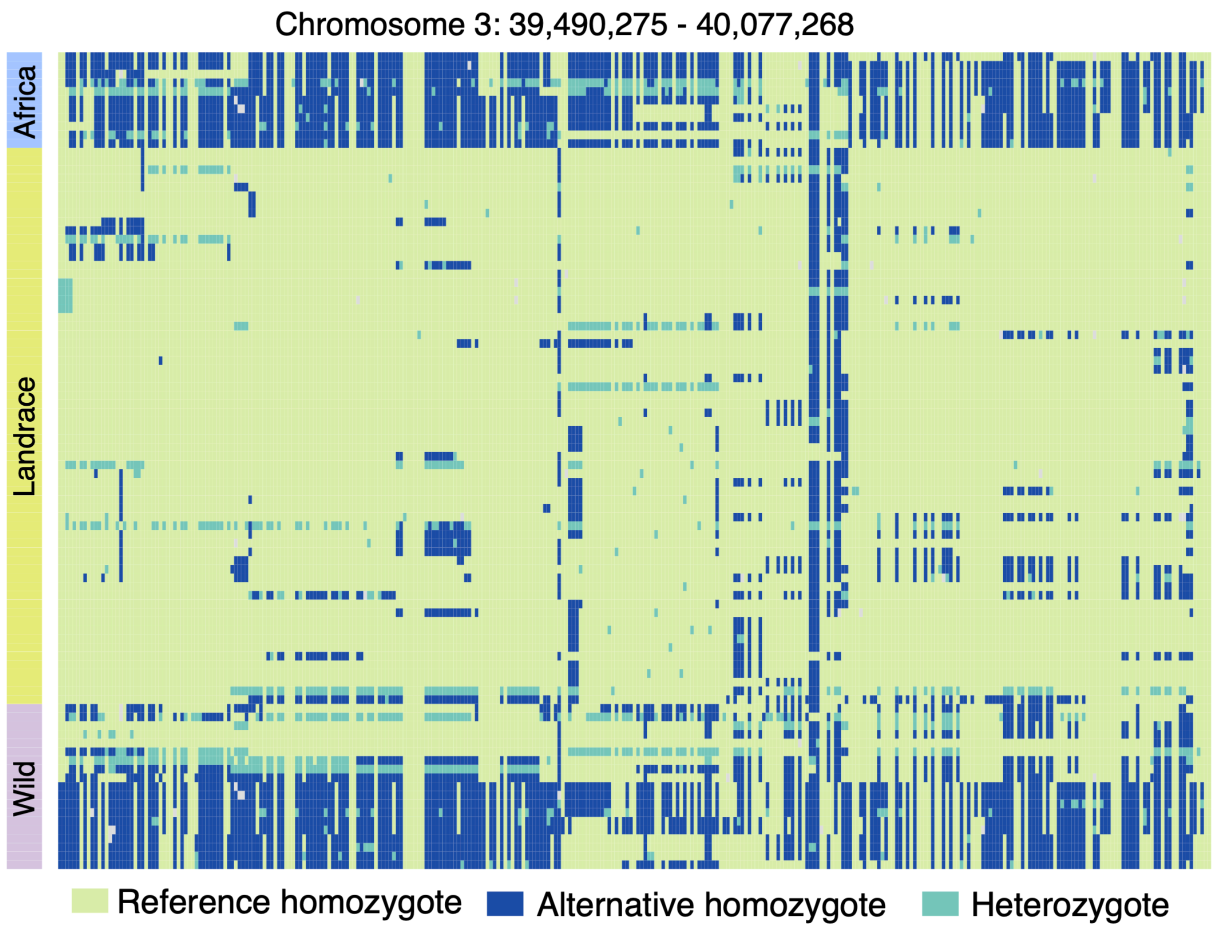


**Supplementary Fig. 12. Heatmap of genotype profiles in an introgressed region from wild to the African population on chromosome 3.**

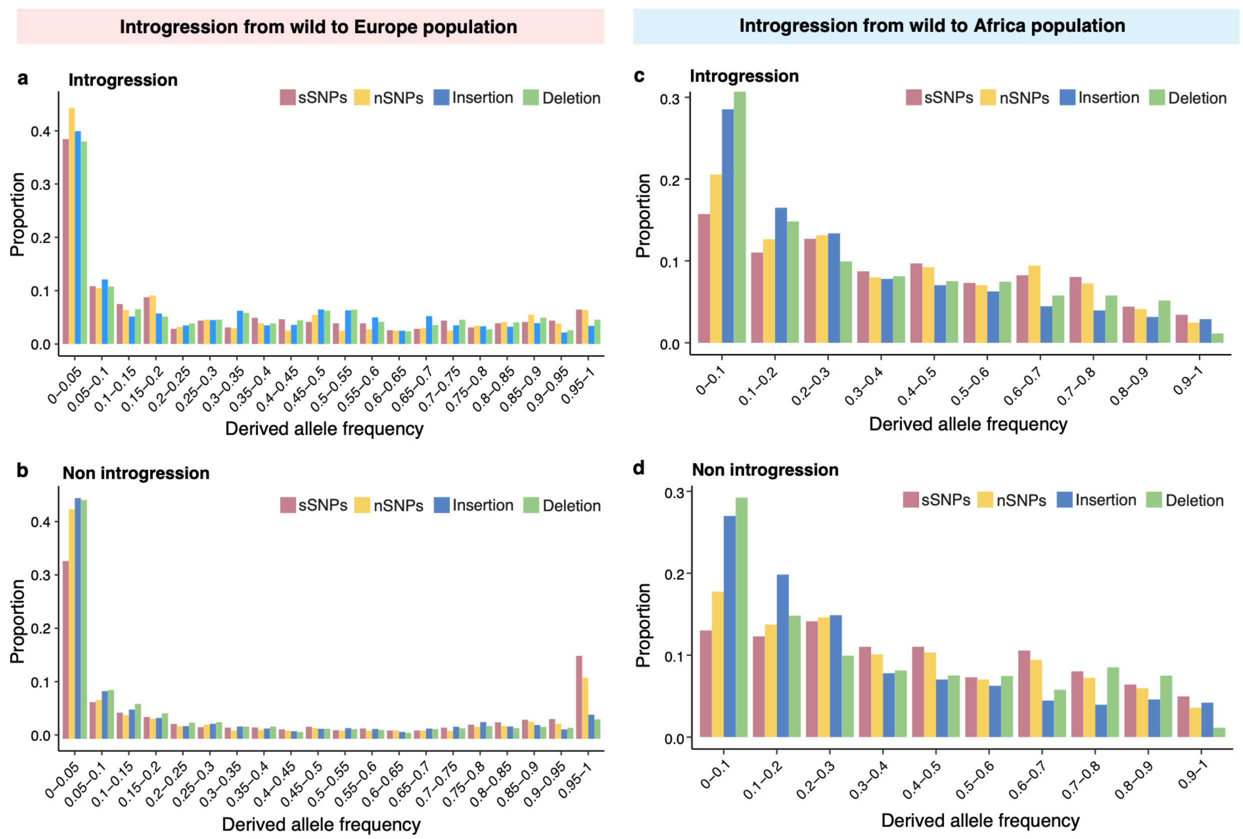


**Supplementary Fig. 13. Site frequency spectrum (SFS) of sSNPs, nSNPs, and SVs in introgressed and non-introgressed regions. a-b,** SFS of sSNPs, nSNPs, insertions, and deletions in regions with (**a**) and without (**b**) introgressions from wild to the European population. **c-d,** SFS of sSNPs, nSNPs, insertions, and deletions in regions with (**c**) and without (**d**) introgressions from wild to the African population.

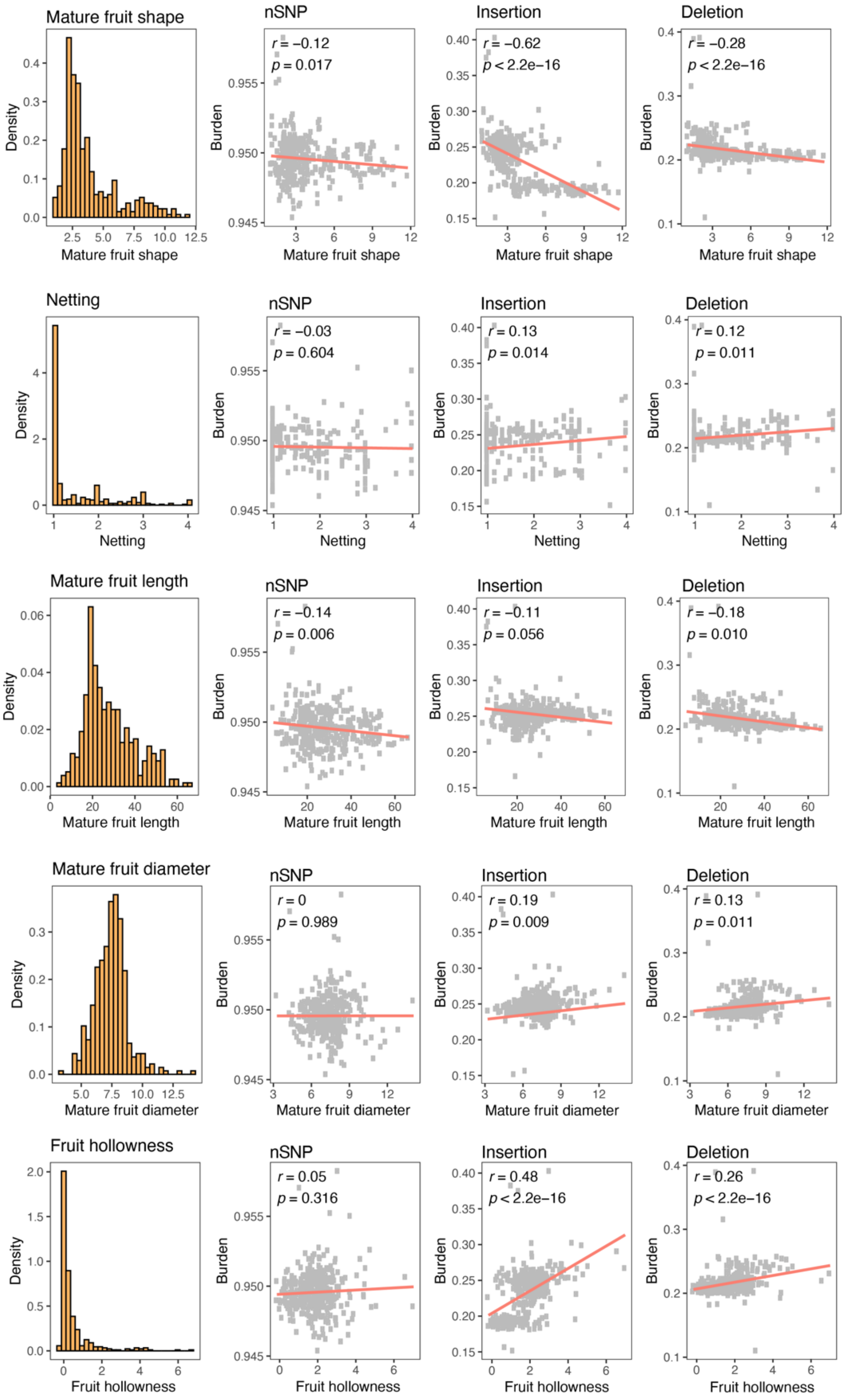


**Supplementary Fig. 14. Correlations between cucumber agronomic traits and SNP/SV burden.** The leftmost panels show the phenotype distributions of cucumber traits. nSNP, non-synonymous SNP.

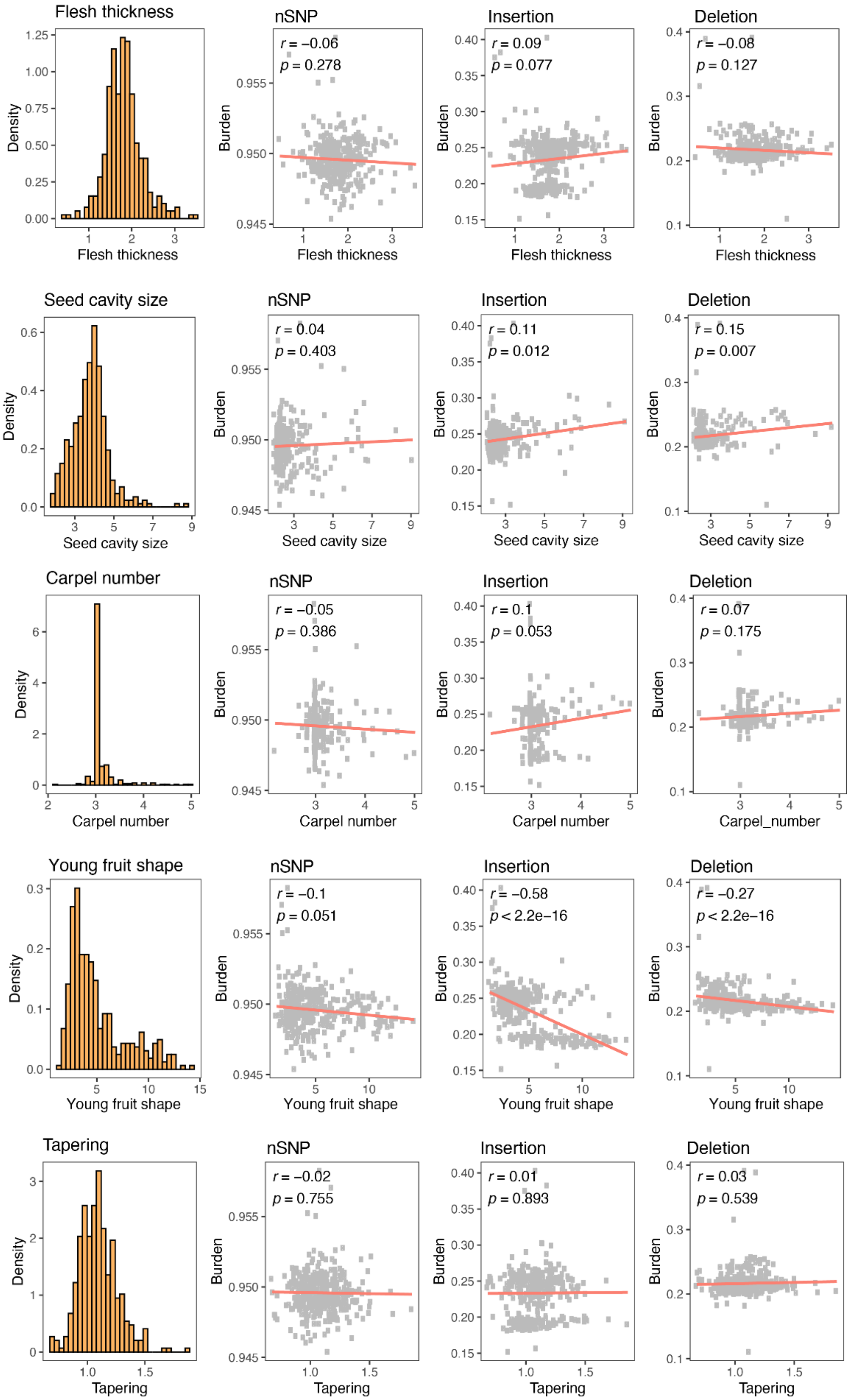


**Supplementary Fig. 14. Continued.**

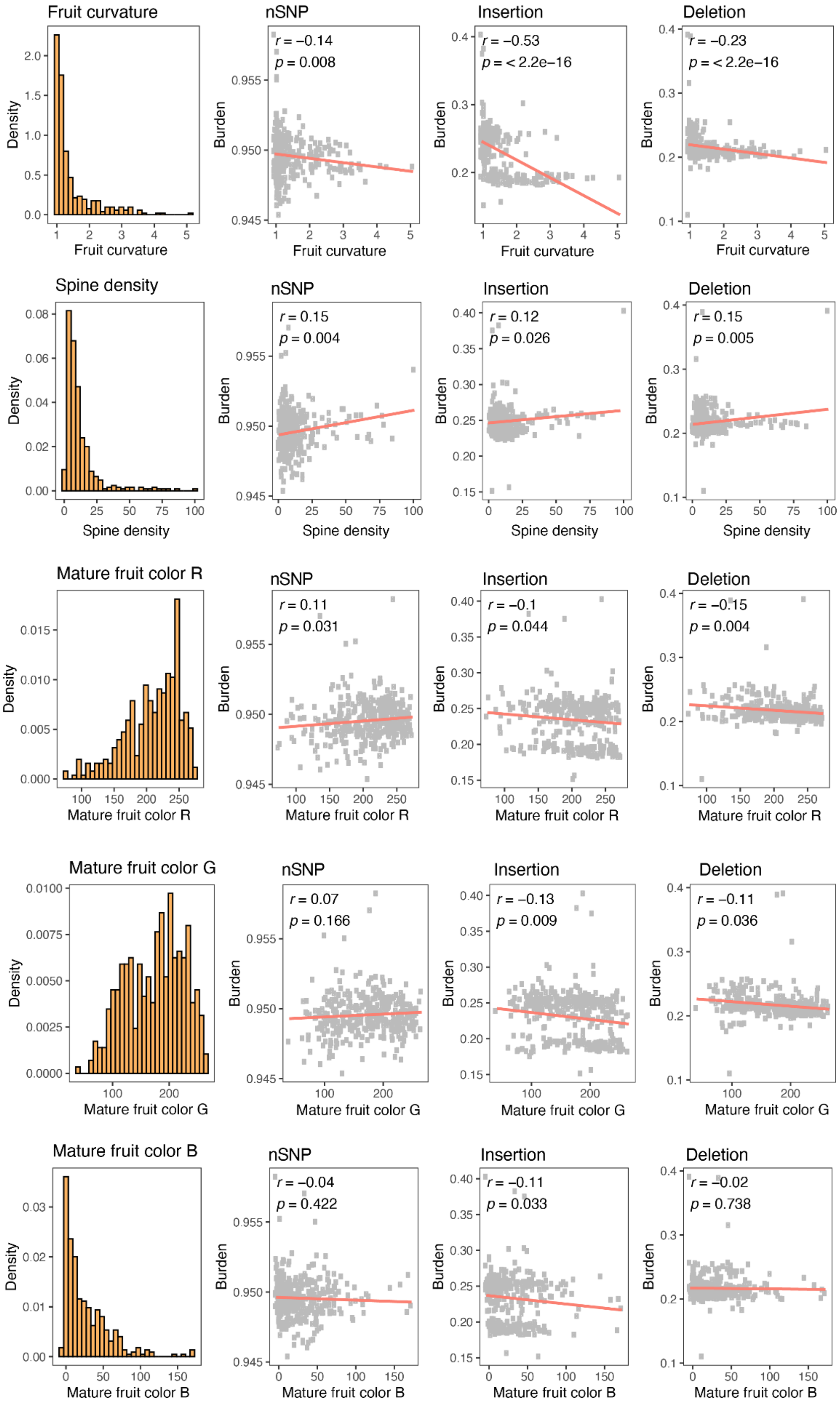


**Supplementary Fig. 14. Continued.**

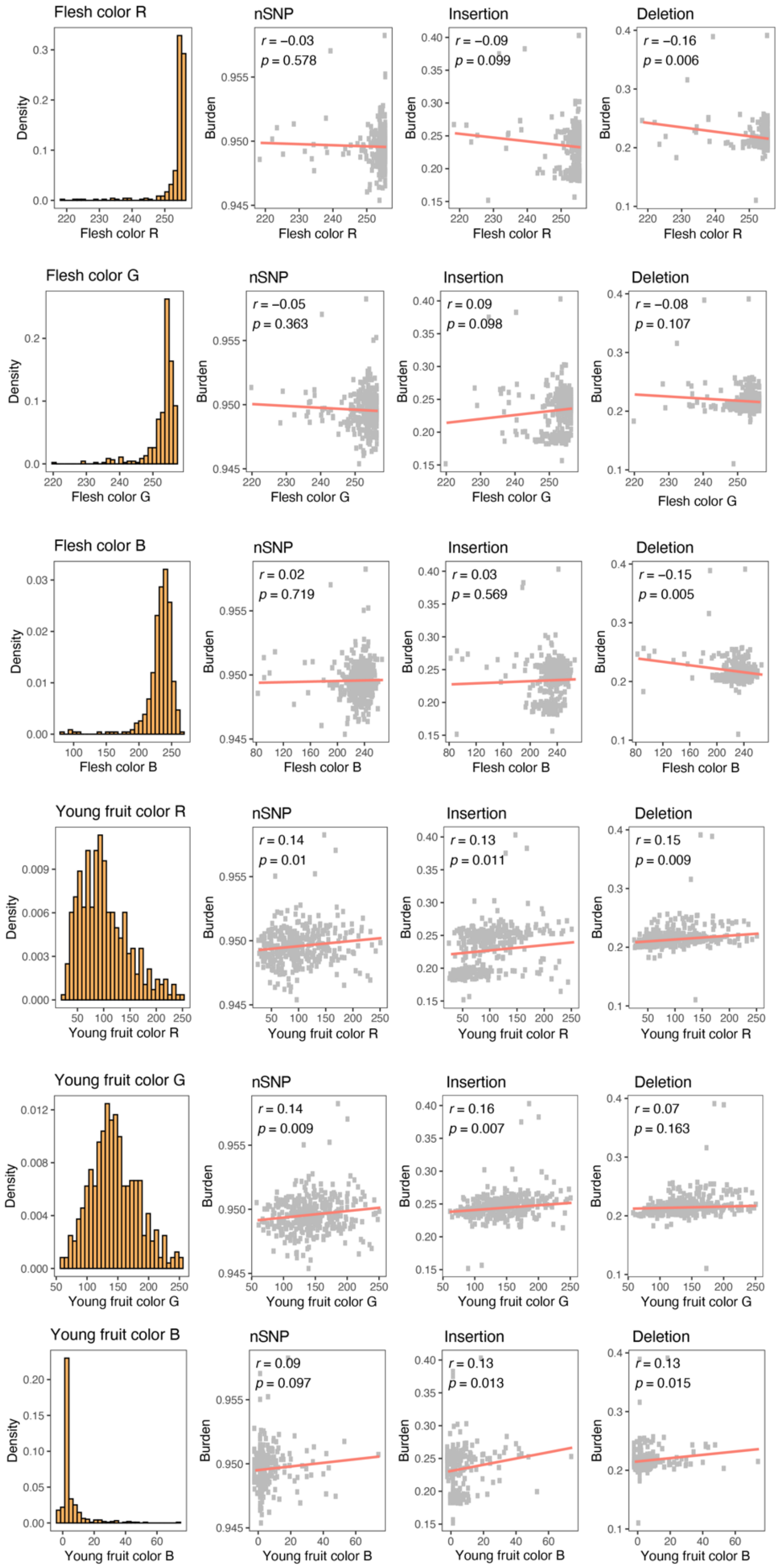


**Supplementary Fig. 14. Continued.**

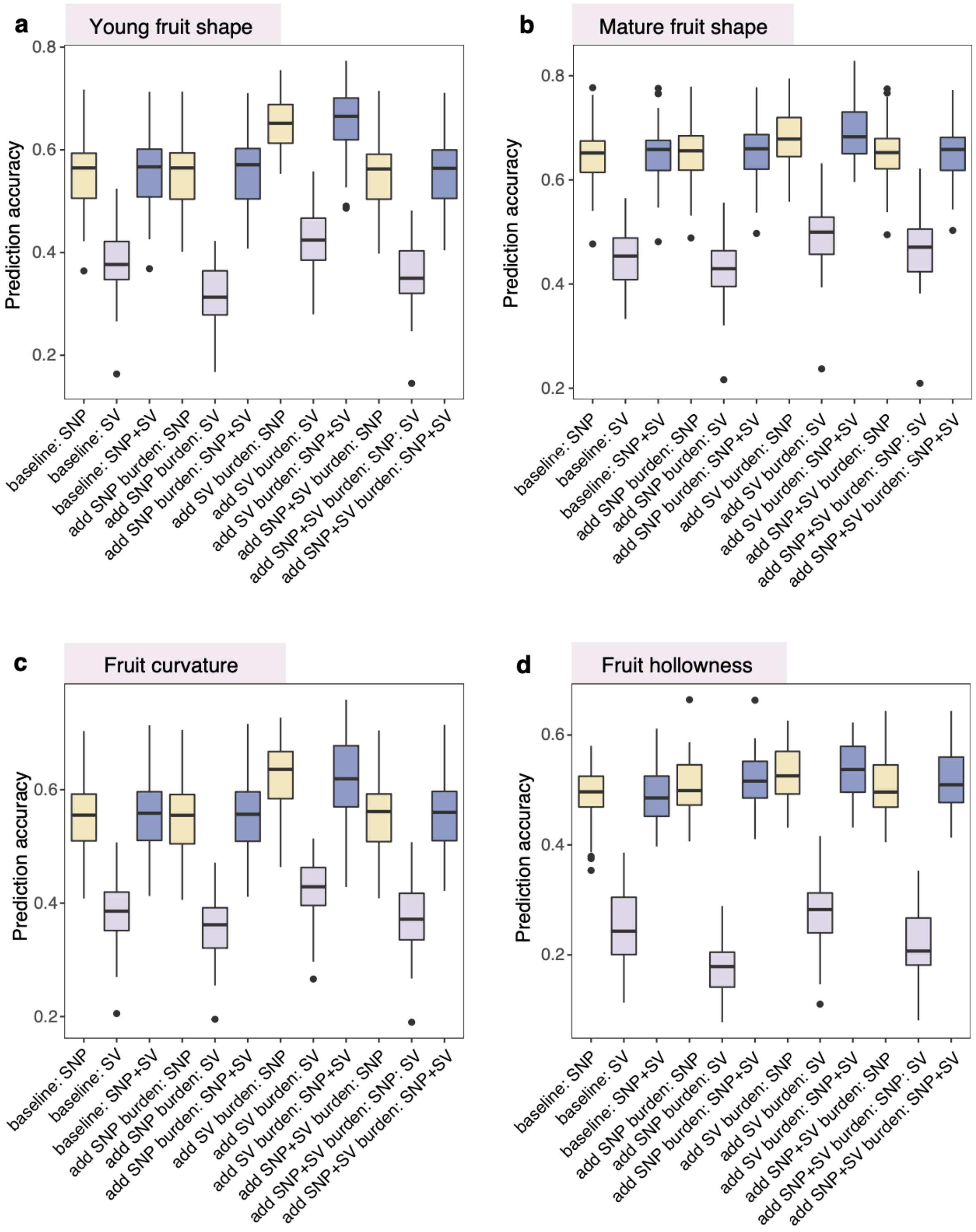


**Supplementary Fig. 15 Genomic prediction accuracies for four traits significantly correlated with SV burden. a-d,** Genomic prediction accuracies for young fruit shape (**a**), mature fruit shape (**b**), fruit curvature (**c**), and fruit hollowness (**d**). For each boxplot, the lower and upper bounds indicate the first and third quartiles, respectively, the center line indicates the median, and the whiskers extend to 1.5× the interquartile range.

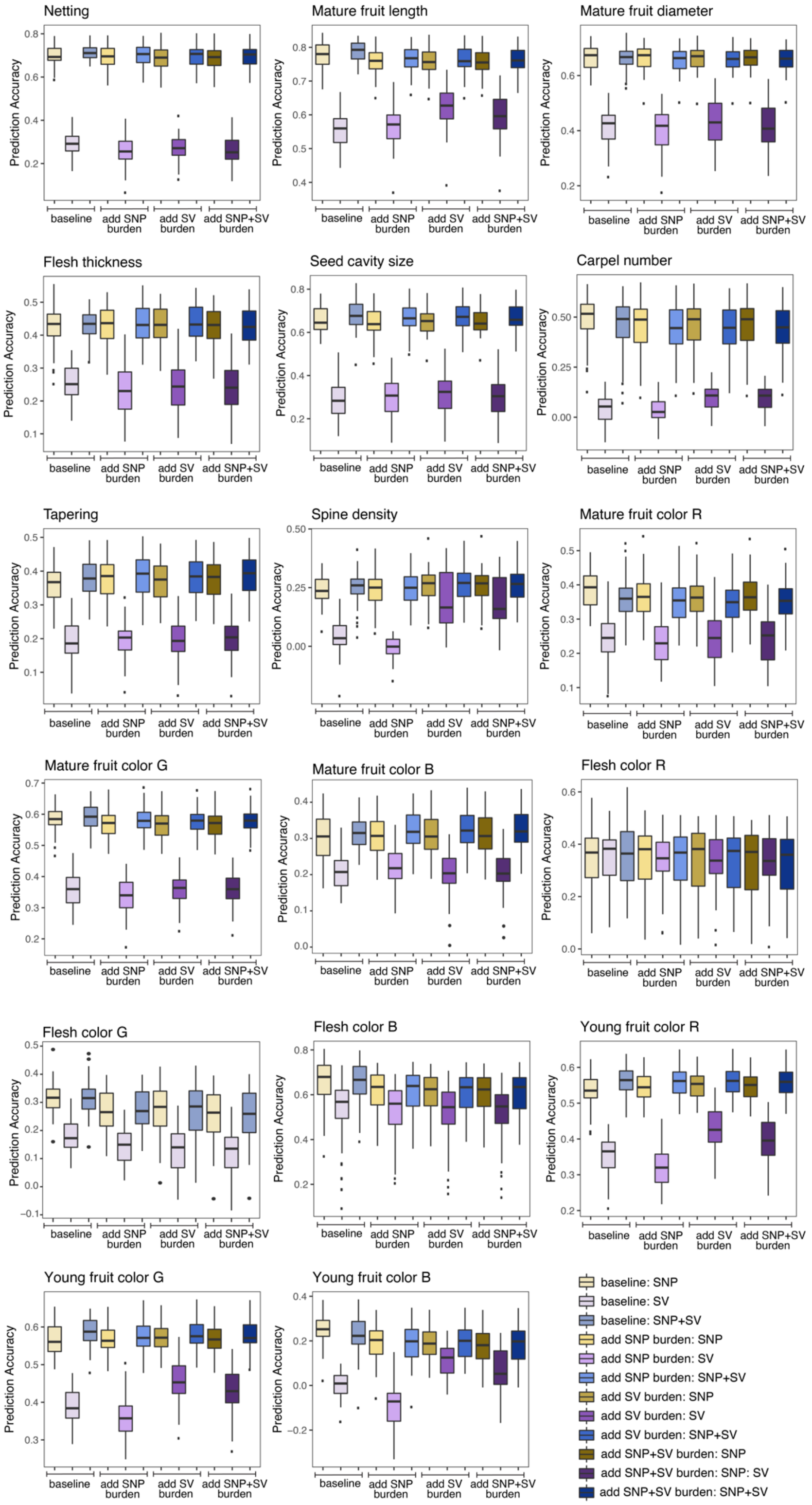


**Supplementary Fig. 16 Genomic prediction accuracies for traits not correlated with SV burden.** For each boxplot, the lower and upper bounds indicate the first and third quartiles, respectively, the center line indicates the median, and the whiskers extend to 1.5× the interquartile range.
